# Cell-Based Sensor for Extracellular DNA

**DOI:** 10.64898/2026.08.19.745795

**Authors:** Boao Xia, Nicholas A. Kalogriopoulos, Ruoxin Wen, Zachary M. Lane, Honghao Li, Nicolas Buitrago, Sangsin Lee, Richard D. Gao, Isaac Ive, Yerim Kim, Alice Y. Ting, Jerzy O. Szablowski

## Abstract

Detection of molecules with cell-based sensors allows for conversion of binding events into gene expression outputs. Here, we present a cell-based sensor that can detect extracellular double-stranded DNA. This sensor is based on an engineered receptor which we call Luminescent Ultrasensitive Nucleic Acid Reporter, or LUNAR. LUNAR is based on a recently developed Programmable Antigen-gated G-protein-coupled Engineered Receptor (PAGER). PAGERs are a genetic fusion of an auto-inhibitory peptide, a protein-binding domain, and a modified kappa opioid receptor. PAGERs are gated by two binding events. First, a protein ligand displaces an intramolecular inhibitor, Arodyn, then a second ligand activates the receptor. By replacing the protein-binding domain with a DNA binding zinc finger protein (ZFP) we could detect extracellular DNA in a dose-dependent fashion. Here, we show that first-generation LUNAR constructs can detect both oligonucleotides and plasmid double-stranded DNA with nanomolar sensitivity in mammalian cells. Future work will focus on improving sensitivity, fold-change, and multiplexing capabilities for sequence-specific DNA detection.

**GRAPHICAL ABSTRACT:** 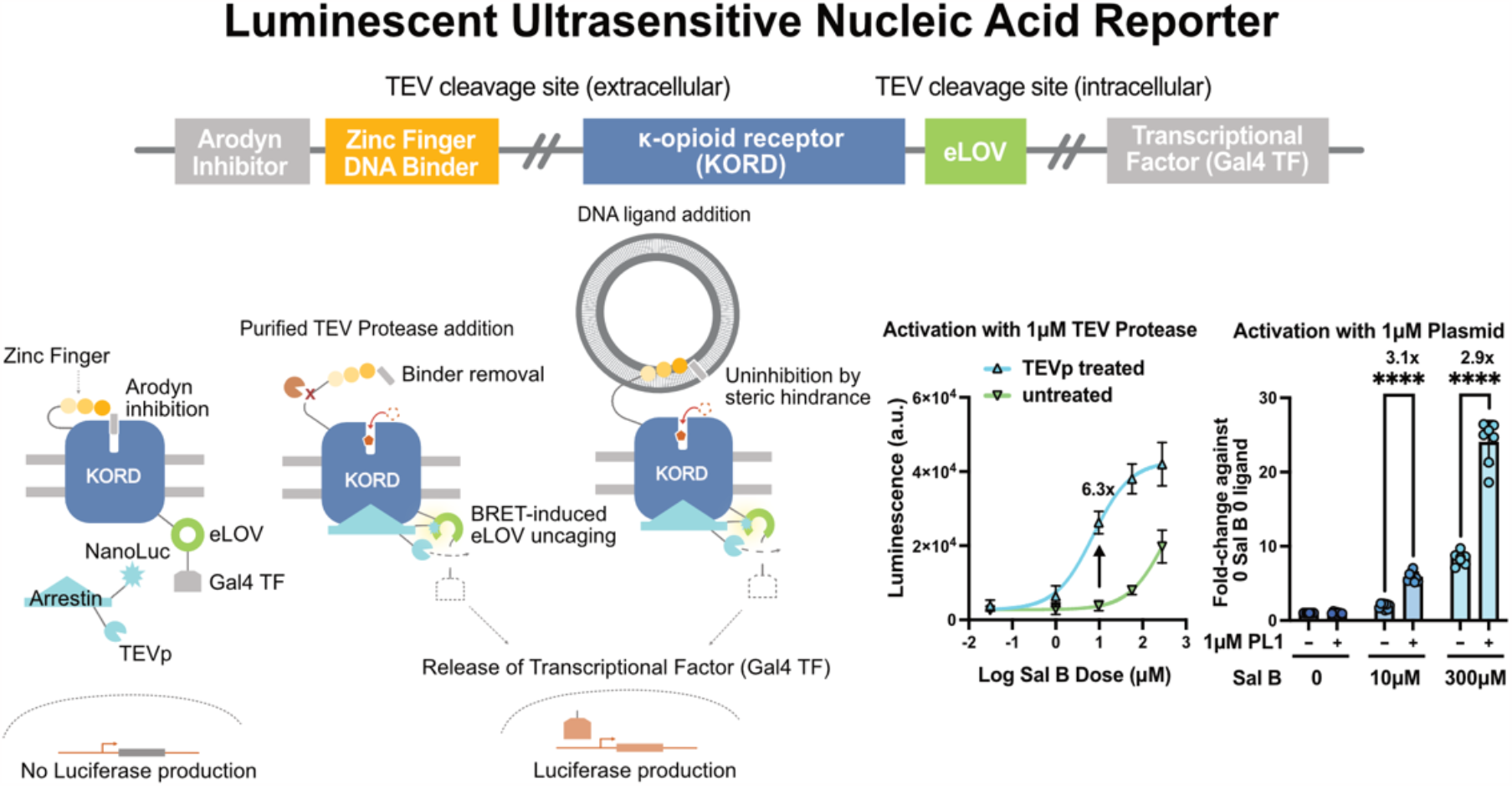

## INTRODUCTION

Understanding and leveraging cellular responses to extracellular biological molecules is a central goal of synthetic biology. To this end, a few molecular tools involving transmembrane signaling have been developed, including Synthetic Notch Receptors (SynNotch)^1^, Modular Extracellular Sensor Architecture (MESA)^2^, and Chimeric Antigen Receptors (CARs)^3^. Recently, a new class of highly modular synthetic receptors has been designed, termed PAGERs, for Programmable Antigen-gated G protein-coupled Engineered Receptors^4^. These receptors bind to cognate antigens via a modular domain on the extracellular side of the membrane. Antigen binding introduces steric hindrance that displaces a peptide inhibitor (Arodyn) of an engineered G-protein coupled receptor (GPCR), such as Kappa Opioid Receptor DREADD (KORD)^5,6^. Once the inhibition is relieved, KORD becomes sensitive to a small molecule agonist Salvinorin B (Sal B) that initiates signaling, ultimately leading to a transcriptional readout, such as the production of firefly luciferase^4,7^. The PAGER construct can be engineered to respond to various soluble antigens, such as EGFP, cytokines and glycoproteins.

Double stranded DNA (dsDNA) is released into bodily fluids under physiologic and pathologic conditions. Sources of dsDNA include mitochondrial DNA (mtDNA)^8,9^, circulating free DNA (cfDNA)^10,11^ and extracellular DNA (eDNA)^12,13^. The level of DNA fragments present in bodily fluids is often associated with various diseases, such as cancer and autoimmune diseases^13–15^. In the brain, dsDNA fragments can be released from hippocampal neurons and glial cells due to energy-intensive molecular adaptations, such as learning^16–18^ and neurodegenerative diseases^19–21^. In addition, environmental DNA detection has been a powerful tool for tracking invasive species in biodiversity and conservation studies^22–24^. While polymerase chain reaction (PCR) based assays^14,25-27^ and Next-Generation Sequencing (NGS)^28,29^ remain the gold standards of dsDNA detection, they require destructive sample processing and *in vitro* signal amplification. As an alternative, cell-based DNA sensors could provide unique advantages such as intracellular signal amplification, or conversion of DNA levels in the extracellular milieu into transcriptional outputs or changes in cellular behavior. However, most cells have no endogenous mechanism to interface with extracellular DNA, meaning they cannot utilize their native intracellular machinery because DNA is generally impermeable to cellular membranes^13^. To address these challenges, we developed Luminescent Ultrasensitive Nucleic Acid Reporter (LUNAR), a PAGER-based receptor capable of detecting dsDNA using living mammalian cells.

## RESULTS

### LUNAR can be activated by the designer drug Salvinorin B (Sal B)

To build LUNAR and allow for extracellular DNA detection, we replaced the protein-binding domain of PAGER with a zinc-finger protein (**ZFP) (Fig. 1a)**. Upon DNA binding, we expected to observe an increase in the luciferase signal. As an initial step, we constructed multiple LUNAR receptors using 3-, 4- or 6-domain ZFPs^30,31^, placed various linkers between the ZFP domains and the peptide inhibitor, and performed circular permutations from within the ZFP domain (summarized in **Fig. 1b, Supplemental Table 1)**. To evaluate the performance of our LUNAR sensors, we transiently transfected a mammalian cell line with plasmids encoding the LUNAR platform and corresponding receptors, then introduced the ligands and analyzed the resulting signals (**Fig. 1c**). We built a library of synthetic 3, 4, and 6 C_2_H_2_ ZF-based LUNAR sensors to investigate the ability of these constructs to sense DNA **(Supplemental Table 1)**.

**Figure 1.**
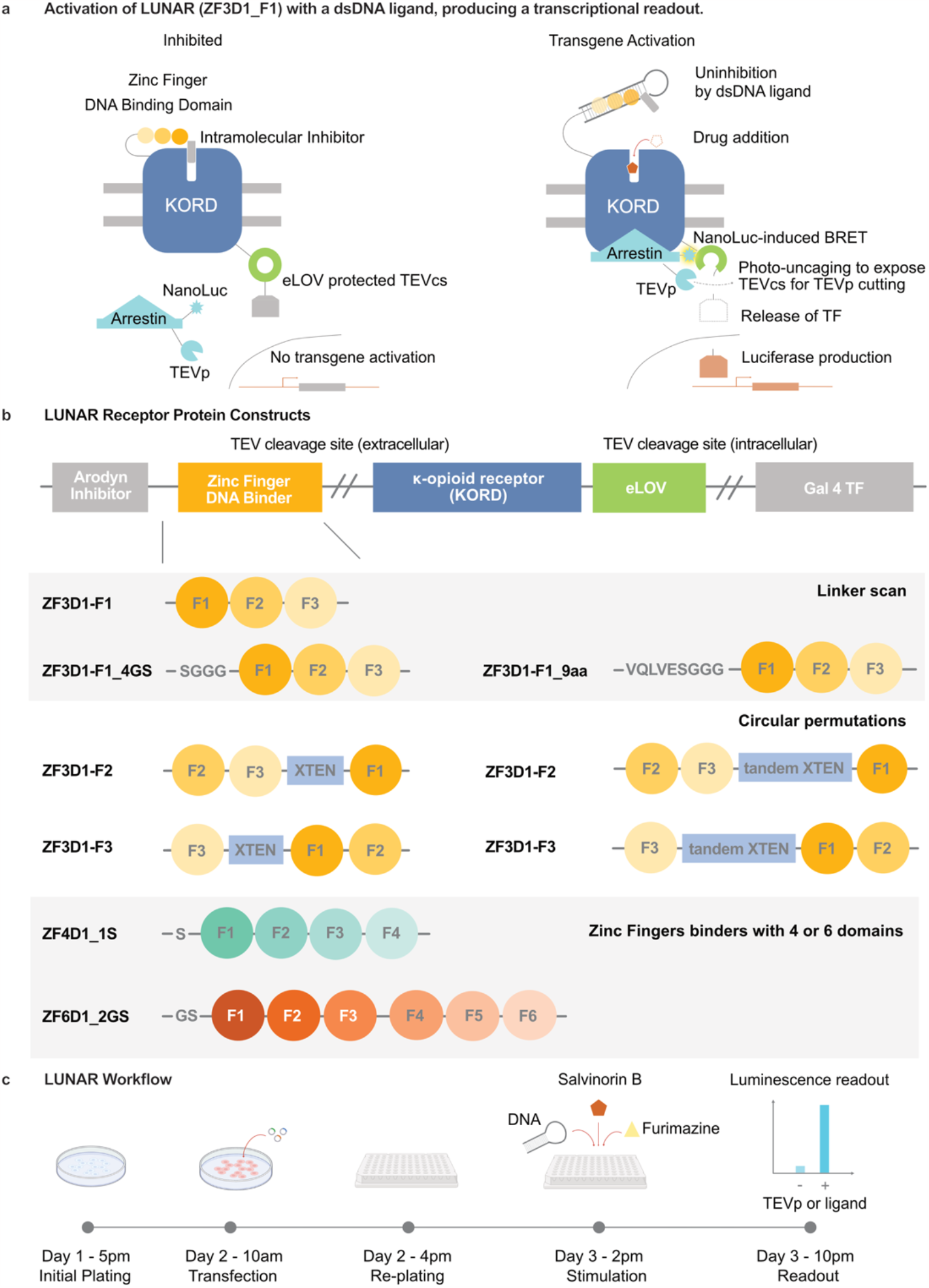
Overview of LUNAR. **a)** Diagram of LUNAR. A κ-opioid DREADD receptor (KORD, blue) is inhibited by the antagonist peptide Arodyn (inhibitor, gray block) fused to its N-terminus. The Zinc Finger DNA binding domain (binder, yellow gradient round patches) is inserted between the inhibitor and KORD such that its binding with the extracellularly supplied annealed dsDNA ligand (ligand, dark gray) forms a complex that sterically interferes with the inhibitor and relieves auto-inhibition, allowing the designer drug Salvinorin B (Sal B, red pentagon) to activate KORD. In the presence of ligand and Sal B and with light activation of the eLOV domain, the NanoLuc-arrestin-TEV protease (NanoLuc-arrestin-TEVp, light blue) is recruited to juxtamembrane, cleaving off the transcription factor Galactose 4 (Gal4, gray) and resulting in the transgene expression of the reporter gene Firefly Luciferase (Luciferase, orange). **b)** Block diagram of the construct of LUNAR, its linker variations, circular permutations, and additional ZF binders built and tested. **c)** Workflow for LUNAR with transcriptional readout.

We first tested the basic functionality of these receptors, such as the ability of each construct to inhibit KORD. Since PAGERs use intramolecular inhibitors to reduce their sensitivity to Sal B, proteolytic removal of such inhibitor should result in improved sensitivity to Sal B. Thus, to confirm the extent of intramolecular inhibition of the LUNAR, we proteolytically cleaved the inhibitor from the extracellular domain of the KORD using Tobacco Etch Virus protease (TEVp, **Fig. 2a)**. We then compared the levels of luciferase signal with and without TEV cleavage at various Sal B doses to quantify the inhibition. A higher fold-change between the two TEV cleavage conditions indicates more effective removal of auto-inhibition by the LUNAR receptors. We evaluated two different illumination methods, external light (light) and Bioluminescence Resonance Energy Transfer (BRET) with a genetically fused NanoLuc. Both illumination protocols increased the firefly luciferase signal, with environmental light providing 3.8 ± 1.2-fold difference compared to no-TEVp control at 10 µM Sal B **(Fig. 2b, 2d)**. The BRET protocol resulted in 6.3 ± 1.2-fold activation at 10 µM Sal B and additionally shifted Sal B’s EC50 from 383.6 nM to 8.4 nM **(Fig. 2c-d)**. Due to the greater fold change, we decided to use BRET for the remaining experiments to evaluate activation of LUNAR with DNA ligands.

**Figure 2.**
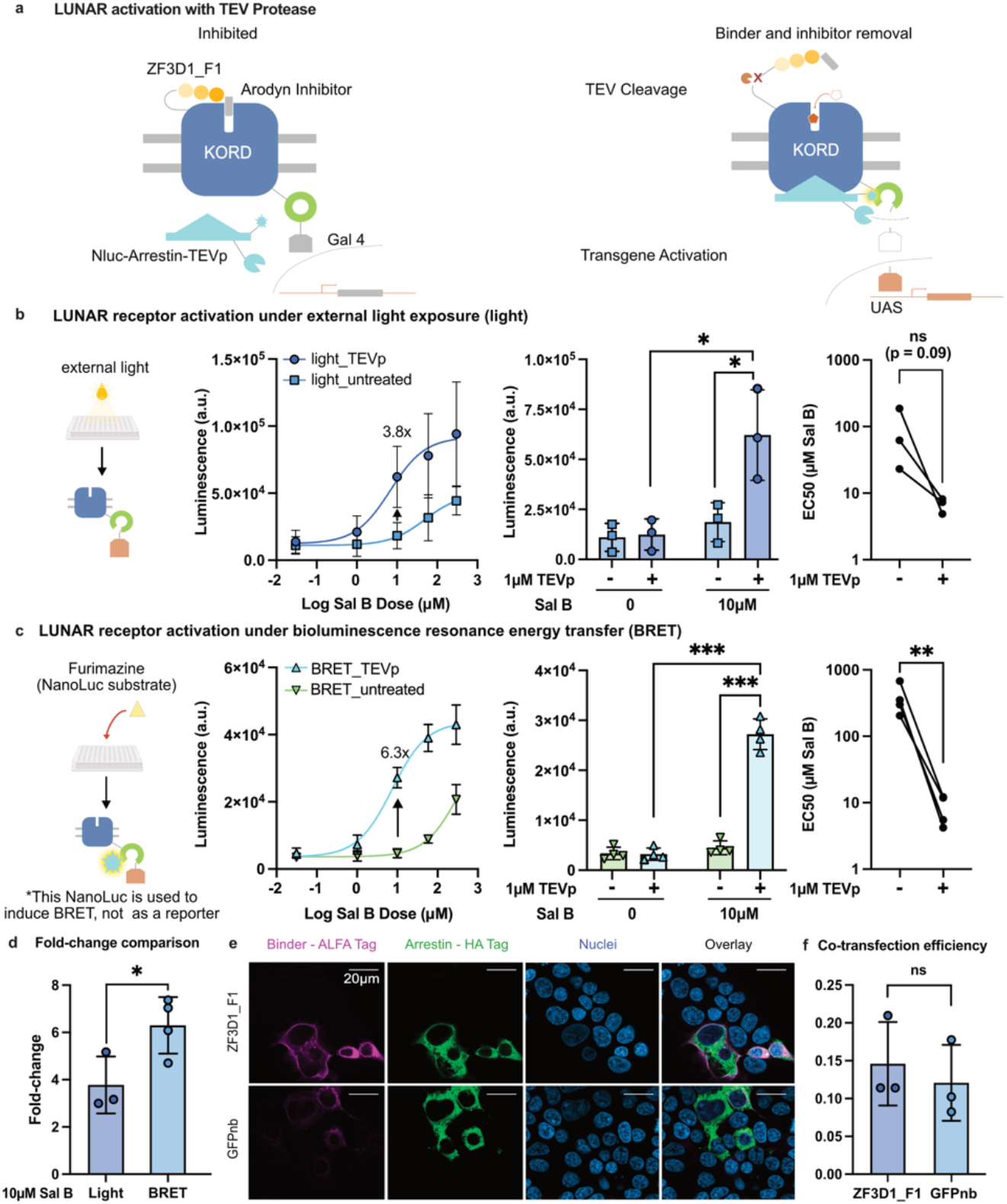
Validation of LUNAR Receptors. **a)** The binder domain can be removed from the receptor by TEV protease, since there is a TEV cleavage site joining the binder and KORD. Removal of the inhibitor allows for KORD activation and release of Gal4 and its binding to the promoter (UAS), which results in transcriptional output. **b)** and **c)** Characterization of receptor responses to Sal B without (-) and with (+) TEV protease (TEVp) treatment under two different illumination for uncaging mechanisms, the bar charts featuring responses at 0 µM and 10 µM Sal B, and EC50 changes (n = 3-4 independent biological replicates, each with 2 technical replicates, *p<0.05, **p<0.01, ***p<0.001, ns = not significant). **d)** Comparison of maximum fold change between two uncaging methods at 10µM Sal B. **e)** Immunocytochemistry showing binder-ALFA Tag localization of the extracellular domain, the localization of Nluc-arrestin-HA Tag-TEVp component at the juxtamembrane, along with the nuclei stain of LUNAR-expressing HEK293T cells. **f)** Quantification of transfection efficiency of both the binder and the arrestin component, showing a co-transfection efficiency of 0.15 ± 0.06 for ZF3D1_F1 and 0.12 ± 0.05 for GFPnb PAGER (3 independent biological replicates, each with 6 field of views, ns = not significant).

For receptor ZF3D1_F1, we confirmed the membrane localization of the receptor and the cytosolic localization of the arrestin component. To achieve this, we performed immunostaining with anti-ALFA tag antibody and anti-HA tag antibody **(Fig. 2e)**, respectively. The ALFA tag is a 13-amino-acid synthetic peptide designed as an epitope tag for protein localization detection, similar to the HA or FLAG tags^32^. We also quantified the fraction of the HEK293T cells expressing both the binder and the arrestin component for ZF3D1_F1 LUNAR (0.15 ± 0.06) and compared it against GFPnb PAGER (0.12 ± 0.05) and found no significant difference between the two **(Fig. 2f)**.

### Linker Length affects LUNAR activation

To evaluate the effects of the ZFP structure on DNA binding in the context of LUNAR we tested two linker variants between the DNA-binding domains of a three-domain ZFP without circular permutations (**Fig. 1c**, ZF3D1_F1). The first ZFP was fused directly to the peptide inhibitor while the others had 4- and 9-amino acid extensions (ZF3D1_4aa, ZF3D1_9aa, respectively). The flexible linkers were introduced to optimize the relative positioning of the peptide inhibitor with respect to the binding pocket of the DREADD to allow for maximum inhibition. One variant included “4aa linker”, with a SGGG sequence^33^, and the other, “9aa linker”, with a VQLVESGGG sequence. The former was derived from GGGGS, which is broadly used for domain separation while the latter was also used in the GFP PAGER providing a plausible spatial separation of the inhibitor and the molecule-binding domain of LUNAR^4^ **(Supplemental Fig. 1a)**. The Sal B activation with BRET light exposure of these constructs showed that linker extension reduced the magnitude of activation. LUNAR without the extended linkers (ZF3D1_F1) showed the highest fold activation of 6.3 ± 1.2-fold at 10 µM Sal B, compared to 4.3 ± 1.4-fold by the 4aa linker variant at 60 µM Sal B, and 1.6 ± 0.2-fold by the 9aa linker variant at 60 µM Sal B **(Supplemental Fig. 1b-c)**. The ZF3D1-F1 also had the highest magnitude of SalB EC50 shift between TEVp treated and untreated conditions, suggesting the most efficient intramolecular inhibition **(Supplemental Fig. 1d)**. We have also modeled the inhibitor-binder-transmembrane domain in AlphaFold3^34^ with the addition of Zinc ions as co-factors, and Arodyn inhibitor placed within the orthosteric pocket of KORD for the ZF3D1_F1 without any linkers in top 5 predicted structures **(Supplemental Fig. 2a)**, adding the linkers results in only 1 out of 5 top predicted structures showing the intended placement of Arodyn **(Supplemental Fig. 2b-c)**. Consequently, we chose to evaluate the ZF3D1-F1 construct in greater detail.

### Influence of the DNA ligand length and structure on LUNAR activation

The size and structure of DNA ligands in LUNAR will likely determine the efficacy of intramolecular inhibitor displacement, receptor activity, and SalB binding. To evaluate the overall effects of various DNA ligands on the LUNAR activation, we designed a series of constructs **(Fig. 3a)**. First, we made a single target hairpin ligand (SH) – a short 11 bp ds-DNA region containing the 9bp consensus binding sequence to the ZF3D-F1 and a 6nt Adenosine hairpin **(Fig. 3a**, SH). The hairpin design was chosen to avoid the possibility of strand dissociation. Second, we designed a short dsDNA ligand with the same consensus sequencing but no hairpin region **(Fig. 3a**, SL), and finally a longer linear ligand with four repeats of the consensus sequence of ZF3D1 to evaluate whether the longer length and multiple binding sites can improve DNA-dependent LUNAR activation **(Fig. 3a**, LL) and a circular plasmid ligand **(Fig. 3a**, PL1). To predict the effective dose-range of ligand activation, we used a fixed Sal B concentration (EC20, 10 µM) with varying concentrations of the DNA ligands to find the ligand response range **(Fig. 3b)**. We found that gradient SH concentrations can generate a maximum fold activation of 3.1 ± 0.5 and predicted EC50 of 54 nM DNA, gradient SL can generate a maximum fold activation of 2.8 ± 0.6 and predicted EC50 of 10.3 nM DNA, gradient LL had a maximum fold activation of 3.3 ± 1.1 and predicted EC50 of 0.36 nM, and PL1 a maximum fold activation of 5.6 ± 1.1 and predicted EC50 of 28 nM, when using 0 µM Sal B and 0 nM ligand as the normalization baseline **(Fig. 3b)**.

**Figure 3.**
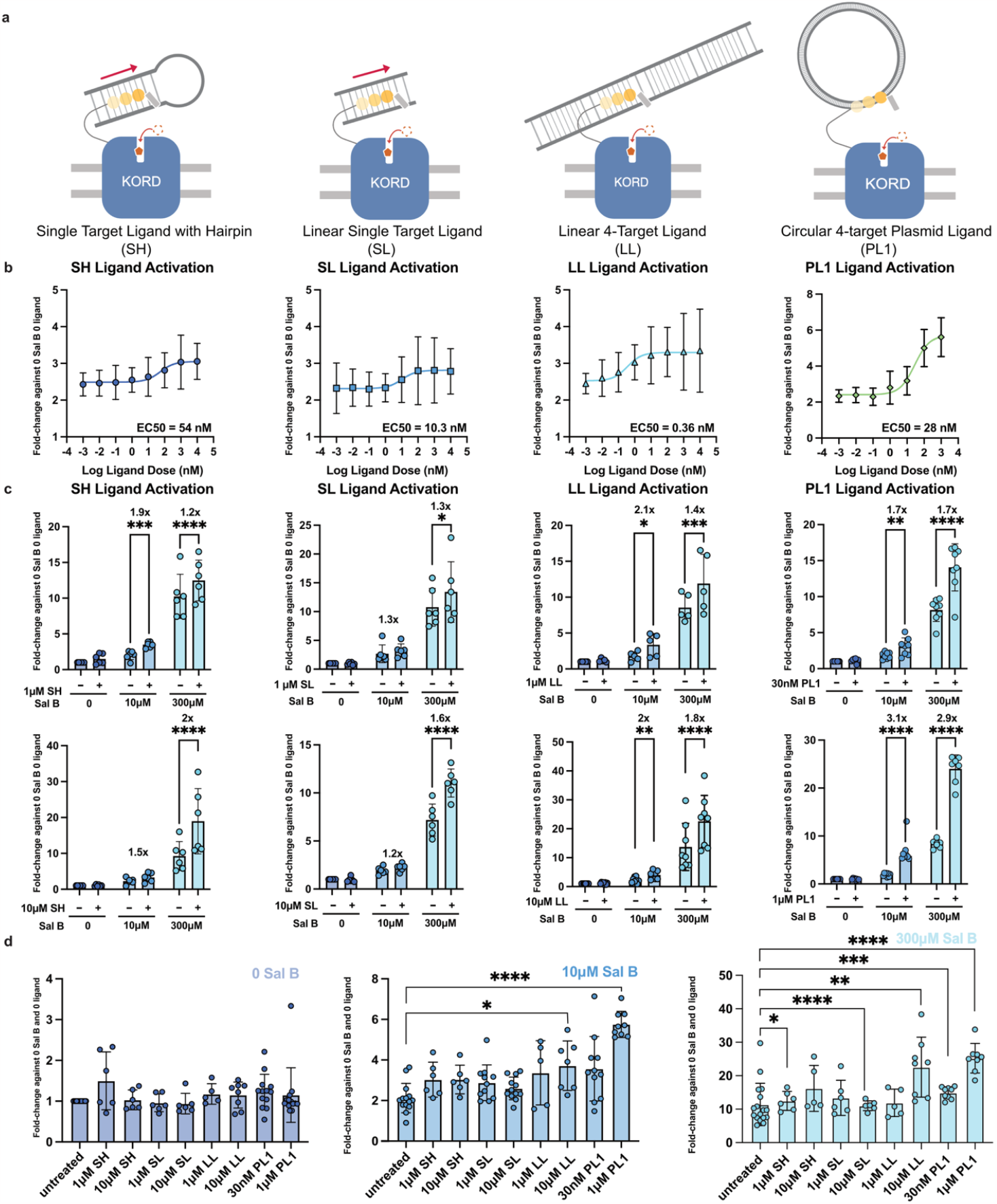
Optimization of DNA Ligands for LUNAR. **a)** Diagram of the DNA ligands used to activate a 3-finger Zinc Finger LUNAR (ZF3D1_F1), from left to right: single target ligand with hairpin (SH), linear single target ligand (SL), linear 4-target ligand (LL), and circular 4-target ligand (PL1). **b)** ZF3D1_F1 was activated with 10 μM Sal B and gradient SH with an EC50 of 54 nM, gradient SL with an EC50 of 10.3 nM, gradient LL with an EC50 of 0.36 nM, and PL1 with an EC50 of 27.8 nM, all using 0 µM Sal B + 0 nM Ligand as the normalization baseline (n = 5-8 biological replicates, each with 2 technical replicates, 2-way ANOVA). **c)** Activation of ZF3D1_F1 with four different DNA ligands and at 30 nM, 1 μM, or 10 μM ligand concentration at 0 µM, 10 μM, and 300 μM Sal B (n = 5-8 biological replicates, each with 2 technical replicates. * = p<0.05, **p<0.01, ***p<0.001, ****p<0.0001). **d)** Comparison of various DNA ligand activation at 0 µM, 10 µM, and 300μM Sal B showing significant activation of DNA ligands against the untreated group. At 0 µM Sal B, none of the ligands led to significant increase of luciferase. At 10 μM Sal B, 1 μM SH, 10 μM LL and 1 μM PL1 obtained significant activation, while 1 μM PL1 gave the highest fold difference of 6.4-fold. At 10 μM Sal B, all DNA ligands except for 1 μM SL and 1 μM LL could be detected, despite fold-change against the untreated group at the same Sal B level diminishing (n = 5-9 independent biological replicates, each with 2 technical replicates, 1-way ANOVA)

We next evaluated the DNA ligand-induced LUNAR activation at several doses of Sal B. We dosed the Sal B to cells cultured in 96-wells at 0 µM, 10 µM, or 300 µM with and without 30 nM, 1 µM or 10 µM double DNA ligands **(Fig. 3c)**. We found that the both ligands with four repeating targets (LL and PL1) showed higher fold change than the single-target ligands (SH and SL), as shown in **Fig. 3c**. Between the single target ligands, SH was more effective than SL at stopping inhibition at 1 µM, suggesting the ligand’s structure can affect the displacement of intramolecular inhibitor of LUNAR. Between the four-target ligands, the circular plasmid PL activated signaling more consistently and with a higher fold change **(Fig. 3c)**. For each individual ligand, the higher concentration group tend to result in a higher and more robust activation of the receptor **(Fig. 3c)**.

We then compared all activation fold-changes at three selected Sal B conditions: 0 µM, 10 µM and 300 µM. We found that at 0 µM Sal B, DNA ligands elicited no activation; at higher Sal B doses we observed significant activation with 10 µM LL and 1 µM PL1 compared to the untreated condition at 10 µM Sal B. We also observed activation at 1 µM SH, 10 µM SL, 10 µM LL, 30 nM PL1 and 1 µM PL1 at 300 µM Sal B **(Fig. 3d)**.

We have also predicted the structures of the LUNAR receptors bound to their DNA ligands using AlphaFold3^34^. All five models of the four dsDNA ligands (SH, SL, LL, and a section of PL1 with four binding targets) show successful displacement of the inhibitor from the orthosteric pocket of KORD **(Supplemental Fig. 3a-d)**. The PL1 ligand contained only a section of the PL1 due to the computational complexity of the entire plasmid.

### Confirmation of molecular specificity of LUNAR ligand detection

To determine specificity of DNA activation of LUNAR (ZF3D1_F1), we prepared stimulation solutions with a protein ligand (EGFP) at different doses. We observed no activation at selected concentrations similar to **Fig. 3b** and **Fig. 3c (Supplemental Fig. 4a)**. As a positive control of GFP-PAGER we compared its activation by EGFP, SH, and PL ligands, and found that only the former led to significant activation **(Supplemental Fig. 4b)**. We therefore concluded that DNA does not activate PAGER design non-specifically in the absence of fused ZFP, and EGFP does not cross-activate LUNAR.

### Circular permutation changes LUNAR sensitivity to Sal B and DNA ligands

We hypothesized that the orientation of DNA ligand within the ZFP can affect LUNAR activation **(Fig. 4a)** To test this hypothesis, we generated variants of LUNARs in which the DNA binding domains were circularly permuted. Specifically, we changed the order of the individual zinc-finger domains within the ZFP domain of LUNAR and inserted XTEN linkers to provide sufficient flexibility as the ZF DBD fold back into the conformation that binds to DNA targets. Overall, we constructed four variants with the second finger or the third finger of the three-finger binder ZF3D1 leading, and each with single or tandem XTEN linker insertions **(Fig. 4b)**. We named the variants after the leading finger_linker. For example, for the variant with the second finger leading the binder domain, joined by a single XTEN linker, was labeled F2_XTEN. From the TEVp cleavage experiment, we saw that F2_XTEN gave a maximum fold change 4.3 ± 0.4 at 60 μM Sal B, F2_tandem XTEN gave a maximum fold activation of 2.3 ± 0.3 at 10 μM Sal B, F3_XTEN gave a maximum fold activation of 4.1 ± 0.9 at 10 μM Sal, and F3_tandem XTEN gave a maximum fold activation of 5.1 ± 1.5 at 60 μM Sal B **(Fig. 4c)**.

**Figure 4.**
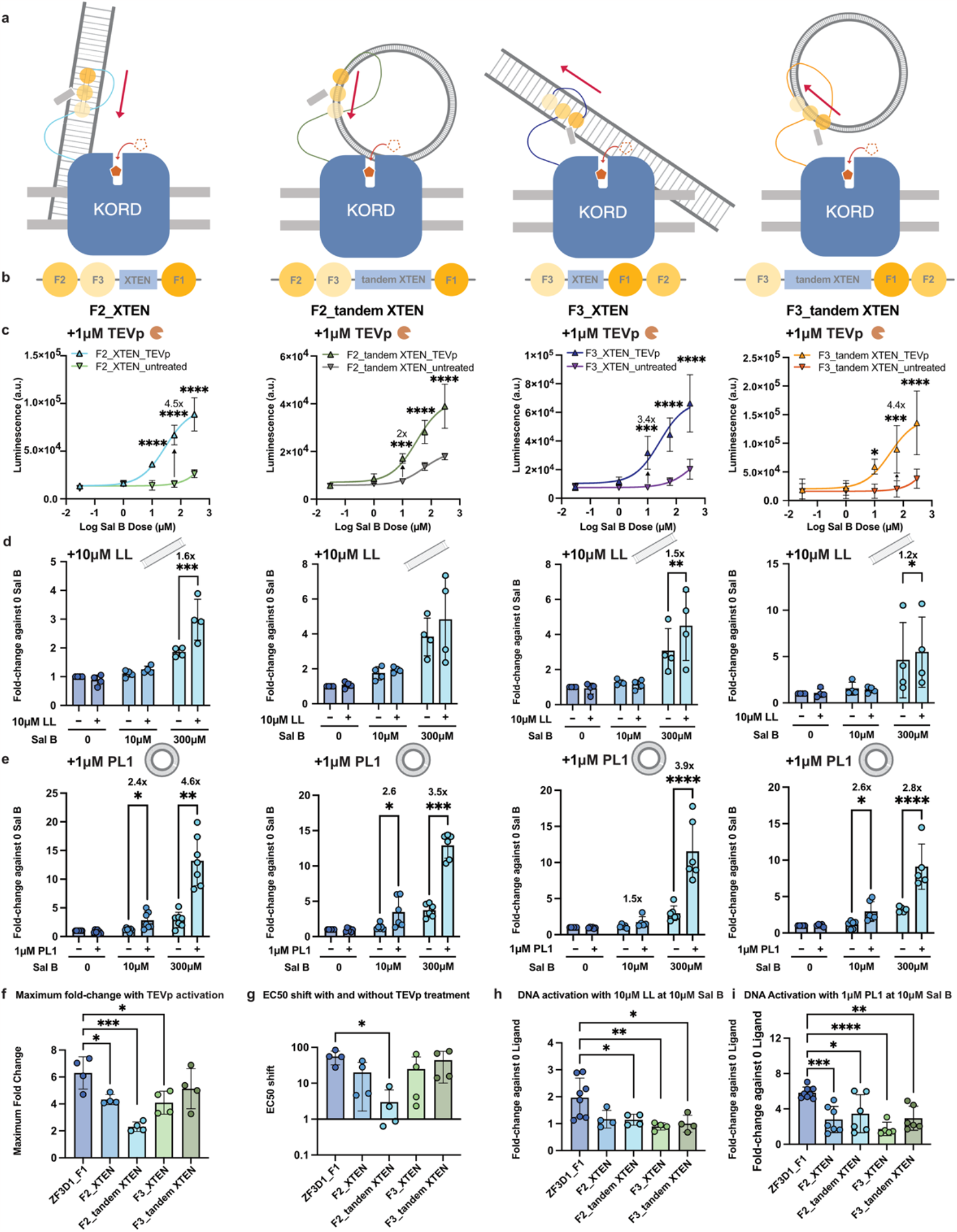
Effect of circular permutation on receptor response and ligand detection. **a)** Diagrams of circular permutations of ZF3D1 3-finger LUNAR receptors being activated by long linear ligand LL or circular plasmid ligand PL1. **b)** Naming of four circular permutations, each with either the second or third finger directly fused to the inhibitor, and containing either single or tandem XTEN linkers. **c)** All four variants were activated by a gradient of Salvinorin B, as characterized by the TEV protease treatment assay. **d-e)** All four variants were activated by long linear ligand LL **(d)** or circular plasmid ligand PL1 **(e)** at selected Sal B concentrations. **f-i)** Comparison of **(f)** maximum fold change, **(g)** EC50 shift, and **(h)** fold change against 0 nM ligand at 10 μM Sal B for both 10 μM LL and **(i)** 1 μM PL1 across five circular permutation variants (original and 4 permutants).(n = 5-8 biological replicates, each with 2 technical replicates, 2-way ANOVA, *p<0.05, **p<0.01, ***p<0.001, ****p<0.0001).

We then dosed the two 4-target DNA ligands to all four permutation configurations and found that they can be activated by 4-target linear (LL) and circular (PL) DNA ligands **(Fig. 4d-e)**, though to a lesser extent than the original conformation **(Fig. 3c)**. From the results, F2_XTEN showed significant sensitivity at 300 μM Sal B for both ligands, while also a significant fold-increase at 10 μM Sal B for PL1; F2_tandem XTEN showed significant activation at 300 μM Sal B for 1 μM PL1 only; F3_XTEN and F3_tandem XTEN showed significant activation by both ligands at 300 μM Sal B, but not by PL1 at 10 μM Sal B (p<0.05; 2-way ANOVA). When the fold changes were compared, we found significant differences in their fold-activation **(Fig. 4f)** and EC50 shift **(Fig. 4g)** and the significant difference in fold-change by the long linear ligand LL was dosed **(Fig. 4h)** and comparable fold-change by the plasmid ligand across the variants **(Fig. 4i)** at 10μM Sal B (1-way Anova). Comparisons of maximum fold change **(Fig. 4c)** and EC50 shift **(Fig. 4d)** from the TEVp cleavage experiment, as well as the fold-change caused by DNA ligand from across the permutants, suggested that, in the case of this ZF3D1, circular permutation might compromise the DNA binding capacity **(Fig. 4f-i)**.

The AlphaFold3 modeling showed successful binding of Arodyn to the KORD’s binding pocket F2_XTEN in some of the top models **(Supplemental Fig. 5a**, 3 out of 5 models), F2_tandem XTEN **(Supplemental Fig. 5b**, 3 out of 5 models), and F3_XTEN **(Supplemental Fig. 5c**, 2 out of 5 models), and failed inhibition in F3_tandem XTEN **(Supplemental Fig. 5d**, 0 out of 5 models). The lower rate of inhibition could lead to higher background signal, and therefore could lower fold-change compared to ZF3D1_F1, which is consistent with the data in **Fig. 4f-g**.

We fed the inhibitor-ZFP-KORD domain and linear DNA structures (LL or a 135bp section of PL1 with four binding sites) as a complex to Alphafold3, and obtained the predicted structures of the binders and their relative position to the ligands. While both DNA ligands displaced the inhibitor in all four permutants in at least some of the models, the addition of DNA induced some structural changes to the ZF binders. To quantify the structure distortion of ZF domains, we used the perimeter of the triangle formed by the three Zn ions from ZF3D1_F1 as a baseline. The perimeter is 92.7 ± 0.6 Å with the LL ligand and 92.1 ± 0.4 Å with the fraction of the PL1 ligand. In the LL ligand addition groups, for F2_XTEN, 4 out of 5 models maintained ZF conformation **(Supplemental Fig. 6a)** with a perimeter of the Zn triangle of 92.7 ± 0.6 Å, while the final stretched model had a perimeter of 116.8 Å; for F2_tandem XTEN, 3 out of 5 models had tighter ZF domains with a perimeter of 90.2 ± 8.6 Å, while 2 had stretched ZF domains with a perimeter of 142.7 ± 32.3 Å due to stretched DNA binding sites **(Supplemental Fig. 6b)**. For F3_XTEN, 3 out of 5 models maintained tight ZF domains with a perimeter of 94.9 ± 9.1 Å **(Supplemental Fig. 6c)**, and the 2 stretched cases had a perimeter of 122.1 ± 5.4 Å. For F3_tandem XTEN, 2 out of 5 models maintained tight ZF domains with a perimeter of 74.4 ± 4.2 Å **(Supplemental Fig. 6c)**, while the stretched cases resulted in a perimeter of 106.2 ± 43.2 Å. In the PL1 additions groups, all models of F2_XTEN and F2_tandem XTEN had ZF domains **(Supplemental Fig. 7a-b)** with F2_XTEN having a Zn triangle perimeter of 75.1 ± 9.8 Å and F2_tandem XTEN having a perimeter of 71.9 ± 3.6 Å. In 2 out of 5 models, F3_XTEN showed tight ZF domains with a perimeter of 94 ± 0.1 Å while the 3 stretched cases showed a perimeter of 126 ± 5.6 Å. For F3_tandem XTEN, the 3 models with tight domains had a perimeter of 93.4 ± 25.7 Å while the 2 stretched cases showed a perimeter of 151.2 ± 38.3 Å **(Supplemental Fig. 7c-d)**. These structure and binding results suggest a higher variation of zinc finger domain positions introduced by the XTEN linker between various receptor constructs. The inhibitor-ZF domains, when separated from the rest of the receptor by XTEN linkers, could reduce the steric clash of the DNA and ZFP with the receptor and arodyn leading to KORD inhibition in the presence of DNA ligand, which could explain the lower sensitivity in the LUNAR assay **(Fig. 4h-i)**.

### Four and Six Finger LUNAR can be activated by TEVp and plasmid DNA ligands

To understand the effect of ZFP binding sequence length, we thought to design binders comprised of longer DNA-binding domains. Specifically, we used 4- and 6-finger zinc finger DNA binding domains referred to as ZF4D1_1S and ZF6D1_2GS, respectively **(Fig. 5a)**.

**Figure 5.**
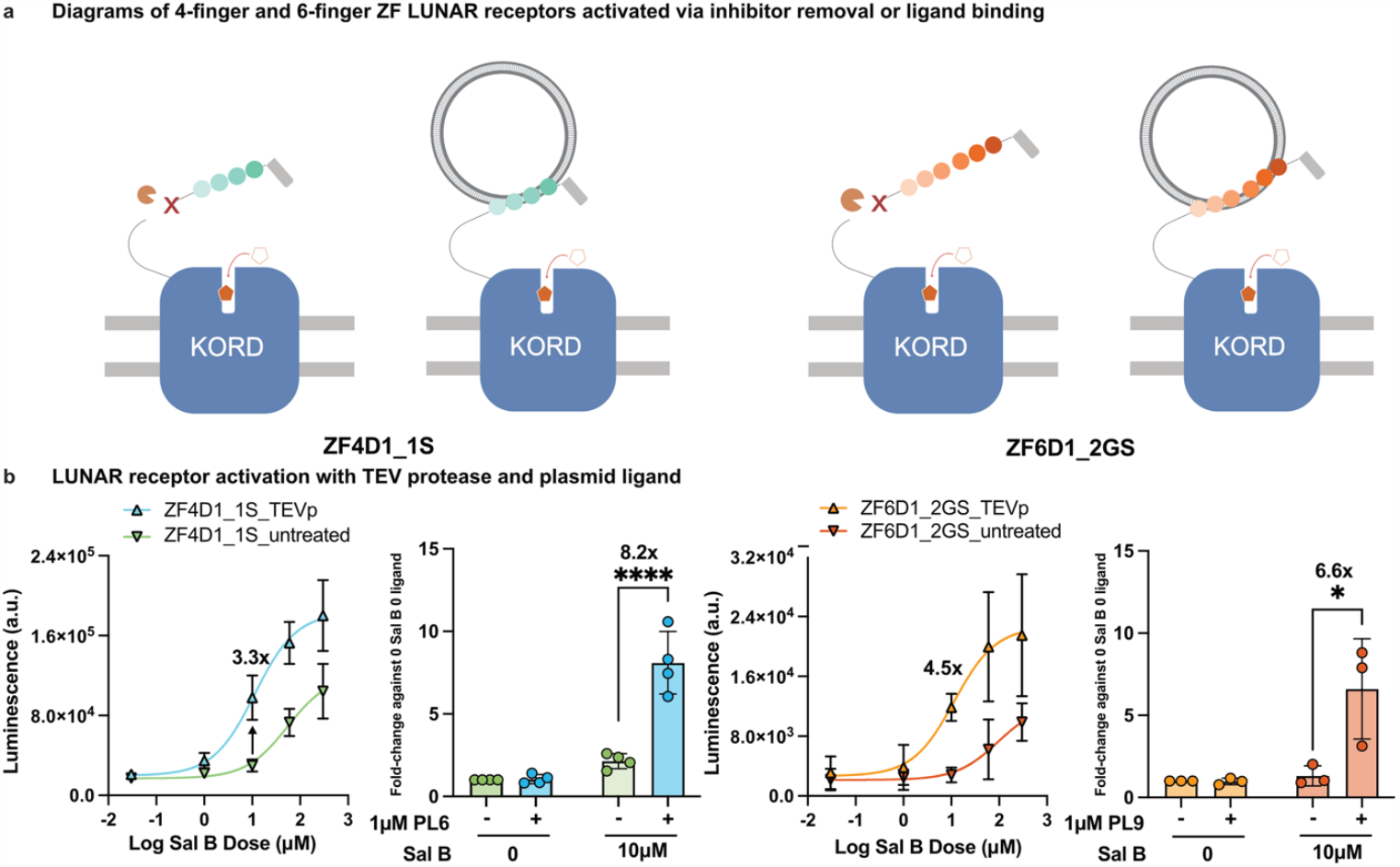
Sal B and DNA activation of four and six finger LUNARs. **a)** Diagram of synthetic receptors fused with a 4-finger ZF DNA binding domain or a 6-finger ZF DNA binding domain being activated by externally supplied TEV protease or plasmid ligand. **b)** Dose response curves of gradient Sal B on ZF4D1_1S and ZF6D1_2GS LUNAR receptors show activation via either inhibitor removal or ligand binding similar to ZF3D1_F1 LUNAR. ZF4D1_1S has a maximum fold activation of 3.3 at 10 μM Sal B with an EC50 shift of 4.9 from without (-) to with (+) TEVp, and ZF6D1_2GS has a maximum fold activation of 4.5 at 10 μM Sal B with an EC50 shift of 8.3 between TEVp treatment groups. Both multi-finger LUNARs can also be activated by their respective 4-target circular plasmid DNA ligands, PL6 and PL9. And both showed significant activation 10μM Sal B by 1μM plasmid DNA. (n = 3-4 biological replicates, each with 2 technical replicates, * = p<0.05, **** = p<0.0001).

We confirmed the default inhibition and efficient removal of the intramolecular inhibitor with a Sal B dose response test and TEVp addition **(Fig. 5b)**. We found that both ZF4D1_1S and ZF6D1_2GS showed sensitization of Sal B activation when the Arodyn inhibitor was proteolytically removed **(Fig. 5b)**. The four-ZFP domain ZF4D1_1S showed 3.3 ± 0.6-fold activation at 10 μM Sal B with an EC50 shift of 4.9 ± 0.8-fold from without (-) to with (+) TEVp. The six-ZFP domain ZF6D1_2GS showed 4.5 ± 1.4-fold activation at 10 μM Sal B with an EC50 shift of 8.3 ± 1.5-fold **(Fig. 5b)**. We also designed and constructed the respective plasmid ligands for ZF4D1(PL6) and ZF6D1(PL9) and reached 8.2 ± 3.3-fold activation with 1 μM PL6 at 10 μM Sal B for ZF4D1 and 6.6 ± 1.1-fold activation with 1 μM PL9 for ZF6D1 at 10 μM Sal B **(Fig. 5b)**.

Similar to the previous evaluation of ZF receptor and receptor-DNA ligand binding for the three-finger ZF3D1, we modeled the inhibitor-ZFP-KORD domain of 4- and 6-finger LUNAR with AlphaFold3. We first included the receptors by themselves, and then with a section of the respective plasmid ligands. We obtained the placement of Arodyn within the binding pocket of Kord in 3 out of 5 predicted structures for ZF4D1_1S **(Supplemental Fig. 8a)** and 5 out of 5 cases showed displacement of the Arodyn inhibitors upon the addition of DNA **(Supplemental Fig. 8b)**. To reduce the computational complexity, we also decided to exclude the transmembrane domain from the model **(Supplemental Fig. 8c)**. We found that in this simplified model, all four fingers were wrapping around the major groove of the double stranded DNA target. This prediction is consistent with the experimental result we obtained from the LUNAR assays, which showed plasmid DNA PL6 effectively displacing the inhibitor through binding to the ZF4D1 Zinc Finger binder **(Fig 5b)**. For ZF6D1_2GS, 0 out of the 5 models showed successful inhibition of KORD by the Arodyn peptide inhibitor **(Supplemental Fig. 9a)**, despite the TEV protease assay providing significant activation **(Fig. 5b)**. All 5 models showed inhibitor displacement **(Supplemental Fig. 9b)** and the simplified ZF-single target modeling showed that all six fingers wrapped around the major groove of dsDNA **(Supplemental Fig. 9c)**. The simplified model with isolated ZF binder indicated successful binding between the ZF6D1_2GS Zinc Fingers and its target DNA ligand, which is proved by the activation in the LUNAR assay **(Figure. 5b)**.

### A focused library of LUNARs covers a broad range of analyte concentrations

Given the successful detection of DNA ligands from several LUNAR variants, we went on to expand the repertoire of available receptors. We built a focused library of LUNARs **(Supplemental Table. 1)**, and tested their sensitivity against Sal B with TEVp **(Supplemental Fig. 10)** and plasmid ligand treatments **(Supplemental Fig. 11)**. Among all the receptors tested for proteolytic activation, the highest fold activation was 9-fold, achieved by ZF3D3 at 60 μM, and the plasmid ligands consistently activated these receptors at 10 μM and 60 μM showing statistically significant activation. We found that the testable DNA ligand concentrations resulted in a significant fold-change from 1.7 to 8.2-fold at 1 μM ligand and 10 μM Sal B, with higher fold-changes observed at 300 μM Sal B.

### LUNAR with alternative transcription factor retains sensitivity to Sal B and ligand

To explore the modularity of the intracellular transgene activation pathway of LUNAR, we replaced the Gal4 transcription factor with a reverse tetracycline-controlled transactivator tTA, and replaced the UAS activation sequence with a doxycycline-inducible Tet-ON promoter tre3G **(Supplemental Fig. 12a)**^35^. ZF3D1_F1 with rtTA2-tre3G shows a fold activation of 4.3 ± 0.3 at 10 μM Sal B with an EC50 shift of 11.0 ± 8.4-fold from without (-) to with (+) TEVp, with a p-value of 0.06 **(Supplemental Fig. 12b)**. This synthetic receptor with the Tet-on platform can also sense 1 μM PL1 at 300 μM Sal B with a fold change of 8.5 ± 2.4 against the untreated group **(Supplemental Fig. 12c)**.

## DISCUSSION

In this study, we constructed a series of PAGER-based receptors with DNA binding domains of Zinc Fingers, which we termed LUNARs. Similar to the previously published PAGER receptors^4^, LUNARs are intramolecularly inhibited by a 6 amino acid peptide, Arodyn, until activated by the designer drug Salvinorin B, as shown in Fig. 1a, 2a, 3a, 4a, and 5a. The action of an inhibitor domain can be relieved with addition of TEV protease **(Fig. 2a-c)**, allowing evaluation of the maximum dynamic range. We then analyzed the dose-dependent curves of DNA ligands at a constant Sal B level **(Fig. 3a-b)** and found all four DNA ligands displacing the inhibitor with EC50s in the nanomolar range, with PL1 providing the highest fold-change. To cross-validate the dose-dependent sensitivity with ligand addition, we dosed the DNA ligands at dosages leading to maximum fold-change at selected Sal B concentrations **(Fig. 3c-d)**. We confirmed that the fold-change with and without DNA addition at 10 μM and 300 μM Sal B was matched with the fold changes within the dosage curve in Fig. 3b and d, respectively.

Overall, our sensors detected soluble dsDNA ligands, both small (11bp) and large (2.4kb), with EC50s in nanomolar range (0.36 nM to 54 nM). The addition of linkers between the Arodyn inhibitor and the ZF DNA binding domain reduced the receptor’s response to Salvinorin B **(Supplemental Fig. 1a-d)**. Circular permutation was intended to position the DNA binding closer to the Arodyn to improve displacement and activation fold changes. However, it did not improve the detection sensitivity **(Fig. 4d-e, 4h-i)**. We also designed and confirmed that a library of Zinc Fingers with 3, 4 and 6 domains can respond to Sal B and plasmid DNA ligands with dose dependent effect **(Fig. 5a-b, Supplemental Fig. 4-5)**. Finally, we also validated an alternative transactivation platform tTA-tre3G **(Supplemental Fig. 6)**, which led to a comparable fold-activation and EC50 shift as the Gal4-UAS circuit. indicating the potential to build a multiplex orthogonal DNA detection platform.

While there are other sequence-targeting DNA-binding proteins available, such as Transcription Activator-like Effectors (TALES) and Leucine Zippers, we chose to use Zinc Fingers as the detection module of our dsDNA sensor. The reasons are the compact structure (e.g. to bind to a 9 bp target, the size of Zinc Finger DNA binding domain is around 300 bp, while the size of TALES is around 900 bp) and modularity^36,37^ of Zinc Fingers compared to TALES, and the molecular specificity in protein-DNA interactions^38–40^ compared to Leucine Zippers^41–43^, as the Leucine Zippers often interact with other proteins at comparable binding affinity as DNA targets^44–46^. Because of the relatively compact size of ZFs, the LUNAR construct can be packaged within AAVs, expanding the potential *in vivo* applications of this platform.

LUNAR offers a promising advancement in DNA sensing technology by enabling cell-based sensing of extracellular DNA. With its nanomolar-level sensitivity, along with the genetic encodability, LUNARs can be applied in basic research, clinical diagnostics, and environmental studies. Future work will explore additional ZF domain and ligand designs to increase the dynamic range and fold-changes of LUNARs, and enabling sequence-specific detection of DNA with LUNARs to allow multiplexed chemogenetic control of cell populations.

## MATERIALS AND METHODS

### Plasmid constructs

The LUNAR receptors for transient expression in HEK293T cells were constructed using the standard Gibson Assembly procedure. The backbone was obtained by digesting the GFP-nanobody (LaG2) PAGER receptor plasmid (gifted by the Ting Lab) with NEB restriction enzymes HindIII-HF and KpnI-HF and incubated with Quick CIP, followed by gel extraction. The Zinc Finger DNA binding domains, in original or domain-shuffled configurations, were either ordered as synthesized gene fragments^30^ from Twist Bioscience or PCR amplified from gifted plasmids from the Bashor Lab^31^. Sequences of the ZF DNA binding domains and their consensus DNA sequences used in this study were listed in Supplementary Figure 1. Ligated plasmids were transformed into NEB stable competent E. Coli. cells and extracted by mini or midiprep (Zymo Research, D4200). Plasmids for the Arrestin component and Firefly Luciferase Reporter were gifts from the Ting lab and used without modifications.

### Protein expression and purification

The protein expression plasmid for EGFP was constructed by subcloning the synthetized gene fragment (Twist Bioscience) encoding EGFP with an N-terminal his-tag into the vector plasmid (pSL4-3-pET28a-6xHis-glucm23-mfc) at the NcoI-HF and BsrGI site via Gibson Assembly. 6xHis-EGFP was then transformed in Shuffle T7 Express competent E. Coli cells and purified by Ni-affinity chromatography. In brief, the transformed cells were grown in Terrific Broth medium at 37°C to an O.D. 600 of ~0.6 before induction with 100 µM IPTG at 18 °C for 20 hours. Harvested cell pellets from 1-liter cultures were resuspended in 25 mL of ice-cold lysis buffer (50 mM sodium phosphate, 300 mM NaCl, 5 mM imidazole, 10% glycerol, pH 8.0) containing 1 mM TCEP and protease inhibitor (GoldBio, GB-108-2). Following sonication, the supernatant was centrifuged at 4,000 x g, 4 °C for 20 min, incubated with Ni-NTA agarose resin (Qiagen, 30210) on ice for 1 h and loaded into the glass chromatography column (Bio-Rad, 7372522). The column was then washed with a lysis buffer with increasing imidazole gradient (10 mM to 30 mM, 25 mL each wash) and eluted with 5 ml elution buffer (250 mM imidazole). The eluent was buffer exchanged into PBS using a PD-10 desalting column, and then concentrated with a Corning® Spin-X® UF 20 mL centrifugal filter unit (cutoff at 30 kD). The size and confluency of concentrated proteins were confirmed by SDS-PAGE, and the amount was quantified with the Pierce BCA protein assay (Thermo Scientific, 23225).

### DNA ligand design and preparation

SH were designed as a single stranded DNA oligo containing the target sequence and its reverse complement of a Zinc Finger joined by a short flexible Poly-A region. The other linear ligands (SL, and LL) were prepared by annealing two complementary sequences containing 1 or 4 consensus targets together. These ligands were ordered as synthetized DNA oligos from IDT or Sigma, resuspended in Nuclease-free Duplex Buffer (IDT, 11-01-03-01) at 100 µM and annealed with a temperature gradient curve on a thermocycler, and successful annealing was confirmed by running a gel electrophoresis.

The plasmid ligand (PL9) containing four ZF target regions (gift from the Bashor Lab) was transformed into NEB Stable E. Coli cells. We inoculated the colonies in a 5mL starter culture in Luria Broth (LB) supplemented with Carbenicillin, then further grew it in four liters of LB supplemented with the same antibiotic at 32 °C, shaking at 220RPM for 22 hours, and harvested the cells for a gigaprep (Zymo Research, D4204) to obtain the mini-CMV driven plasmids. We then performed ethanol precipitation to concentrate plasmid DNA to 3 µM. The other plasmid ligands (PL1 through PL8) were constructed using PL9 as their backbone and gene fragments containing four repeats of the targeted sequences for the zinc fingers as their respective inserts with Gibson Assembly.

Plasmid ligands obtained from gigapreps were concentrated to 10 µM using Ethanol precipitation. In a 50 mL Falcon tube, 5 mL plasmids, 2.5 mL 3 M Sodium Acetate (NaAc), and 17.5 mL ice cold 100% EtOH were added and briefly vortexed to mix. The mixture was then stored at −20 °C overnight. On the second day, the mixture was spun down at 4000 g and 4 °C for 40 minutes in a refrigerated table centrifuge (Beckman Coulter Avanti J-15R). The supernatant was carefully poured out and about 1 mL mixture including the pellet was carefully transferred to a 2 mL microcube. The mixture was then pelleted at 14000 g and 4 °C for 25 minutes in a microcentrifuge (Eppendorf 5424R). The supernatant was pipetted out. The remaining pellet was washed with 70% EtOH, and airdried for 30 minutes. After that, the pellet was resuspended in 600 µL IDT Duplex Buffer, incubated at 60 °C overnight and mixed before use or storage.

Details about the ligands can be found in **Supplemental Table. 1**.

### HEK293T cell culture and transfection

HEK293T cells were purchased from ATCC (ATCC, CRL-3216^™^) and cultured as monolayers in Dulbecco’s modified Eagle Medium (Corning DMEM, 10-013-CV) supplemented with 10% Fetal Bovine Serum (Corning, 35-011-CV) and 1% (v/v) Penicillin-Streptomycin (Gibco, 15070063). The cells were maintained at 37 °C under 5% CO_2_ and used between Passage numbers 5 and 20.

To prepare the transfection reagent, 100 mg PEI Max powder (Polysciences, 24765-100) was stirred to dissolve in 90 mL of MilliQ H_2_O. Concentrated HCl was then added dropwise to reduce the pH of the solution to less than 2.0 while stirring continued. Once PEI was fully dissolved (~1 h), concentrated NaOH was added dropwise to adjust the pH to ~7.0. More MilliQ H_2_O was added to adjust the final volume of the PEI solution to 100 mL. And the neutralized PEI Max solution was filter sterilized, aliquoted, and stored at −20 °C. Working stocks were kept at 4 °C for a month after thawing.

To prepare human fibronectin (HFN) coated plates for the LUNAR experiments, human fibronectin (Sigma, A3890401) was briefly centrifuged and incubated at 37 °C for 30 minutes. It is then diluted 50-fold in Dulbecco’s PBS (Corning DPBS, 21-031-CV) and filter sterilized to make a 20 µg/mL working solution. Then 1 mL or 70 µL of working HFN solution was added to each well of a tissue culture-treated 6-well clear plates (Corning, 3516) or 96-well white µClear plates (Greiner Bio-One, 655098), respectively. Coating media were incubated at room temperature for 4 hours, aspirated and left to dry for 20 minutes, and sealed and stored at 4 °C. The plates were pre-equilibrated in a 37 °C and 5% CO_2_ TC incubator before cell plating.

The DNA transfection complex (for one well of a 6-well plate) were prepared right before transfection on Day 2 of the LUNAR assay. In a 5 mL conical tube, we mixed 420 ng receptor plasmid, 120 ng Arrestin component, and 180 ng UAS-Luciferase with 72 µL blank DMEM, we then added 3.6 µL PEI Max and mixed again. The complex was incubated at room temperature for 20 minutes and diluted to a final volume of 2.5 mL with complete DMEM. To perform transfection, we replaced the media from initial plating entirely with complete DMEM containing the transfection complex, wrapped the whole plate in foil (Thor Labs, BKF12), and incubate the plate at 37 °C under 5% CO_2_ for 5 hours until replating.

### Firefly Luciferase Reporter LUNAR experiment

The LUNAR experiment follows a similar protocol to the PAGER and SPARK^7^ studies. On Day 1, approximately 18 hours before transfection, HEK293T cells were plated at a density of 900,000 cells per cell in a PDL-coated 6-well plate. The plates were incubated at 37 °C and under 5% CO_2_. On Day 2, when the confluency of cells was at ~70-90%, the cells were combined with the transfection complex prepared as detailed above and the whole plate was wrapped with a piece of black foil. After 5 hours of incubation, each well was trypsinized with 2 mL Trypsin-EDTA (Corning, 25050CI), neutralized with 2 mL complete DMEM and centrifuged at 300g for 5 minutes to pellet. The cells were resuspended in 10 mL of complete DMEM (Fisher Scientific, MT10013CV, supplemented with 10% FBS and 1% P/S) and replated in ~100 µL of cell suspension to a white, clear bottom 96-well plate. The plate was wrapped in black foil again to protect it from light and incubated at 37 °C for 22 h to allow continuous gene expression of the LUNAR components.

Stimulation including a TEV protease (TEVp) treatment control group took place in a dark TC hood under red light (which does not induce the conformational change of the eLOV domain). One hour before stimulation, growth media was removed by dabbing media on paper towel, and 50 µL complete DMEM containing 1 µM TEV protease (UC Berkeley, QB3 Marco Lab, TEVp) was added to the positive control wells, while the same amount of regular complete media was added to all other wells. During inhibitor cleavage, stimulation solutions with a final amount of 50 µL per well were prepared containing gradient curves of Salvinorin B (Hello Bio, HB4887, 0 µM to 300 µM from stock solution at 20 mM in DMSO) or gradient curves of DNA ligands (0 µM to 10 µM for linear and hairpin ligands annealed at 100 µM, or 0 µM to 1 µM from for plasmid ligands concentrated to 10 µM) at selected Sal B concentrations. For stimulation, TEVp or control media were removed and 50 µL stimulation solution containing Nano-Glo® Live Cell Substrate (Promega, N2012) was added to each well using a multi-channel pipette. The plate was then incubated in the TC incubator at 37 °C and 5% CO_2_ for 30 minutes. After stimulation, stimulation media was replaced with 100µL complete DMEM and the plates were wrapped and placed in the 37 °C incubator. Eight hours after, media was removed and the cells were washed once with 150 µL DPBS before 50 µL of Bright-Glo firefly luciferase substrate (diluted 1:1 in DPBS, Promega, E2620) was added. Each plate was incubated for 30 seconds and orbitally shaken for 1 minute in a microplate reader (Tecan Infinite M2000 Pro) and luminescence signal was read with 1000 msec integration time.

### Quantification and statistical analysis

For each experiment, a TEVp treated group with Sal B gradient was included as a positive control, and an untreated group with Sal B gradient as a standard. The Sal B or ligand dose response curves were normalized against 0 nM dose DNA ligand and at 0 µM dose Sal B. For each experiment, technical duplicate to triplicate was averaged to one data point, and all the biological replicates were plotted with non-linear regression (curve fitting).

All graphs including statistical results were created using GraphPad Prism 10. Error bars represent standard deviation. For comparison across different stimulation modalities or selected dosage groups, p-values were determined using t-test, one-way ANOVA or two-way ANOVA as appropriate, and significance was defined as **** = p < 0.0001, *** = p < 0.001, ** = p < 0.01, * = p < 0.05, ns = not significant.

### Immunocytochemistry and fluorescence imaging of LUNAR

To verify membrane co-localization of the LUNAR receptor and the Arrestin component, immunocytochemistry and fluorescence imaging were performed. Five hours after transfection, HEK293T cells were plated at HFN coated ibidi Chambered coverslip with 8 wells (ibidi, 80827) at approximately 20,000 cells/well. Forty hours after replating, cells were double rinsed in DPBS, fixed in 4% PFA at room temperature for 10 minutes, and double rinsed again. The fixed cells were permeabilized in 0.5% Tween 20 for 15 minutes, and then blocked in 3% BSA for 1 hour. The immunostaining solution was prepared by diluting both conjugated antibodies anti-Alfa tag-AF647 and anti-HA tag-AF488 at 1:500 in the blocking buffer. 300uL staining solution was added to each well, and the cells were incubated at room temperature in dark for 6 hours and rinsed twice with DPBS. The nucleus staining solution was prepared by diluting the Hoechst 33342 stock solution 1:1000 in the blocking buffer. Finally, the cells were incubated with the nucleus staining solution for 10 minutes and double rinsed with DPBS and stored in DPBS until imaging.

Confocal imaging was performed on a Zeiss LSM800 inverted confocal and Airyscan super-resolution microscope. A 40x water-immersion objective and the following solid-state laser lines were used for various fluorophores and dyes: 405nM (Hoechst - Nucleus), 488nM (HA Tag - Arrestin component), and 640nM (ALFA Tag - LUNAR Receptor). All images were collected and processed using the Zeiss ZEN Blue software.

### Structure Prediction with AlphaFold3

The structure of LUNAR receptor-ligand complexes was generated by feeding the protein sequence (Arodyn-ZFP-KORD), the DNA ligand sequence (either as a single strand or as a sense-anti-sense pair) and Zinc ions with the same number as Zinc Fingers in the respective construct to the AlphaFold server^34^. The generated CIF files (five predictions for each model) were then imported to PyMOL^47^ for visualization and quantification.

## ACKNOWLEDGMENTS

The authors appreciate the discussions regarding Zinc Fingers and DNA ligand optimization with Dr. Xiaodong Cheng, Dr. Gang Bao, Dr. Mingjie Dai, Dr. Zhimin Huang, Dr. Lucas Brown and Adam Yaseen. The ZF6D1, ZF6D2 and PL9 plasmids were a gift from the Bashor Lab at Rice University. This research was supported by the David and Lucile Packard Fellowship for Science and Engineering #2021-73005 (J.O.S), NIH R01MH135934 (A.Y.T), and NIH F32CA257159 (N.A.K.).

## AUTHOR CONTRIBUTIONS

B.X., A.Y.T., N.A.K., J.O.S. conceived and planned the research. B.X., Z.L., H.L., N.B., S.L., I.I., and Y.K. did LUNAR experiments with additional input from N.A.K., R.W., and R.D.G. B.X., N.A.K., A.Y.T., and J.O.S. analyzed data. B.X. and J.O.S. wrote the manuscript with input from all other authors. J.O.S. supervised the research.

