## Supplementary Information for "Cell-Based Sensor for Extracellular DNA"

**List Of Supplementary Materials**

**Supplemental Table 1.** DNA and protein sequence of all Zinc Fingers used in this study and their target sequences. p. 2
**Supplemental Fig. 1.** Linker scan and TEVp activation of Zinc Finger-based LUNAR. p. 3
**Supplemental Fig. 2.** Structure Prediction of ZF3D1 receptors. p. 4
**Supplemental Fig. 3.** Structure Prediction of ZF3D1_F1 and various DNA ligands. p. 5
**Supplemental Fig. 4.** LUNAR activation is molecular specific. p. 6
**Supplemental Fig. 5.** Structure Prediction of ZF3D1 circularly permutated receptors. p. 7
**Supplemental Fig. 6.** Structure Prediction of ZF3D1 permutants with long linear DNA ligand (LL). p. 8
**Supplemental Fig. 7.** Structure Prediction of ZF3D1 permutants with an 135bp section of the plasmid DNA ligand (PL1). p. 9
**Supplemental Fig. 8.** Structure Prediction of ZF4D1_1S receptor and matching ligand (PL6). p. 10
**Supplemental Fig. 9.** Structure Prediction of ZF6D1_2GS receptor and matching ligand (PL9). p. 11
**Supplemental Fig. 10.** Sal B activation of all LUNAR constructs. p. 12
**Supplemental Fig. 11.** Plasmid DNA activation of LUNARs. p. 13
**Supplemental Fig. 12.** Characterization of a LUNAR receptor with a different transactivator. p. 14

| No. | Name of Zinc Finger | Feature | DNA Binding Domain DNA Sequence | Length (bp) | DNA Binding Domain Amino Acid Sequence | Length (AA) | Target DNA sequence |
|---|---|---|---|---|---|---|---|
| 1 | ZF3D1 | 3-finger | tctagacccggggagcgccccttccagtgtcgcatttgcatgcggaacttttcgtcccagacccagctggtgcggcatacccgtactcataccggtgaaaaaccgtttcagtgtcggatctgtatgcgaaatttctcccagtccaccaccctgaagcggcatctacgtacgcacaccggcgagaagccattccaatgccgaatatgcatgcgcaacttcagtcagcggaacaacctgggccggcacctaaaaacccacctgaggggatcc | 270 | SRPGERPFQCRICMRNFSSQTQLVRHTRTHTGEKPFQCRICMRNFSQSTTLKRHLRTHTGEKPFQCRICMRNFSQRNNLGRHLKTHLRGS | 90 | tGAAGCAGCAc |
| 2 | ZF3D2 | 3-finger | tctagacccggggagcgccccttccagtgtcgcatttgcatgcggaacttttcgaccaaccagaagctggaggtgcatacccgtactcataccggtgaaaaaccgtttcagtgtcggatctgtatgcgaaatttctccgtgcggcacaacctgcagcggcatctacgtacgcacaccggcgagaagccattccaatgccgaatatgcatgcgcaacttcagtcagcaccccaacctgacccggcacctaaaaacccacctgaggggatcc | 270 | SRPGERPFQCRICMRNFSTNQKLEVHTRTHTGEKPFQCRICMRNFSVRHNLQRHLRTHTGEKPFQCRICMRNFSQHPNLTRHLKTHLRGS | 90 | aGAAGATGGTg |
| 3 | ZF3D3 | 3-finger | tctagacccggggagcgccccttccagtgtcgcatttgcatgcggaacttttcggcccccctccaagctggaccggcatacccgtactcataccggtgaaaaaccgtttcagtgtcggatctgtatgcgaaatttctccctgggcgagaacctgcggcggcatctacgtacgcacaccggcgagaagccattccaatgccgaatatgcatgcgcaacttcagtgacggcggcaacctgggccggcacctaaaaacccacctgaggggatcc | 270 | SRPGERPFQCRICMRNFSAPSKLDRHTRTHTGEKPFQCRICMRNFSLGENLRRHLRTHTGEKPFQCRICMRNFSDGGNLGRHLKTHLRGS | 90 | gGACGACGGCa |
| 4 | ZF3D4 | 3-finger | tctagacccggggagcgccccttccagtgtcgcatttgcatgcggaacttttcgcaccagcgggacctgatgcggcatacccgtactcataccggtgaaaaaccgtttcagtgtcggatctgtatgcgaaatttctccgtgcggcacaacctgacccggcatctacgtacgcacaccggcgagaagccattccaatgccgaatatgcatgcgcaacttcagtgaccccacctccctgaaccggcacctaaaaacccacctgaggggatcc | 270 | SRPGERPFQCRICMRNFSHQRDLMRHTRTHTGEKPFQCRICMRNFSVRHNLTRHLRTHTGEKPFQCRICMRNFSDPTSLNRHLKTHLRGS | 90 | aGTCGATGCCc |
| 5 | ZF3D5 | 3-finger | cccttccagtgtcgcatttgcatgcggaacttttcgacgcggcagaagcttggcgtgcatacccgtactcataccggtgaaaaaccgtttcagtgtcggatctgtatgcgaaatttctccgtcgcgcacaacttgacccgccatctacgtacgcacaccggcgagaagccattccaatgccgaatatgcatgcgcaacttcagtcagcaccccaacctgacgaggcacctaaaaacccacacaggacagaaggac | 255 | PFQCRICMRNFSTRQKLGVHTRTHTGEKPFQCRICMRNFSVAHNLTRHLRTHTGEKPFQCRICMRNFSQHPNLTRHLKTHTGQKD | 85 | aGAAGATGGTg |
| 6 | ZF3D6 | 3-finger | cccttccagtgtcgcatttgcatgcggaacttttcggcgccgagcaagcttgacaggcatacccgtactcatactggtgaaaaaccgtttcagtgtcggatctgtatgcgaaatttctccgatcgctctaatctgacccgccatctacgtacgcacaccggcgagaagccattccaatgccgaatatgcatgcgcaacttcagtgagggggggaacctgatgaggcacctaaaaacccacacaggacagaaggac | 255 | PFQCRICMRNFSAPSKLDRHTRTHTGEKPFQCRICMRNFSDRSNLTRHLRTHTGEKPFQCRICMRNFSEGGNLMRHLKTHTGQKD | 85 | gGACGACGGCa |
| 7 | ZF4D1 | 4-finger | gggatccgaattcccggggaaaaaccgtatgagtgtgatcactgtggaaaatcctttagccagagctctcatctgaatgtgcacaaaagaactcacaccggggaaaaaccgtataaatgcggccagtgtgggaagttctactcgcaggtctcccacctcacccgccaccagaaaatccacaccggggaaaaaccgtatgaatgcgatcactgtgggaaagccttcagcgtcagctccaacctgaacgtgcacagaagaatccacaccggggaaaaaccgtataagtgccctgattgtgggaagagttttagtcagagttccagcctcattcgccaccagcggacacacaccggtgaaaaa | 360 | GIRIPGEKPYECDHCGKSFSQSSHLNVHKRTHTGEKPYKCGQCGKFYSQVSHLTRHQKIHTGEKPYECDHCGKAFSVSSNLNVHRRIHTGEKPYKCPDCGKSFSQSSSLIRHQRTHTGEK | 120 | tGCAAATGGATGAc |
| 8 | ZF4D2 | 4-finger | gggatccgaattcccggggaaaaaccgtacaggtgtaagtactgcgaccgctccttcagcgactcttcgaacctccagcggcacgtccggaacatccacaccggggaaaaaccgtataaatgcggccagtgtgggaagttctactcgcaggtctcccacctcacccgccaccagaaaatccacaccggggaaaaaccgtttcattgtggttattgtgaaaaatctttttctgttaaagattatttgactaaacatattcgtactcataccggggaaaaaccgtataagtgcatggagtgtgggaaggcttttaaccgcaggtcacacctcacacggcaccagcggattcacaccggtgaaaaa | 360 | GIRIPGEKPYRCKYCDRSFSDSSNLQRHVRNIHTGEKPYKCGQCGKFYSQVSHLTRHQKIHTGEKPFHCGYCEKSFSVKDYLTKHIRTHTGEKPYKCMECGKAFNRRSHLTRHQRIHTGEK | 120 | tGGGTATGGAGACc |
| 9 | ZF6D1 | 6-finger | ctctctcaattgccaggcgaacgaccgtttcaatgccggatatgtatgaggaacttctccgaggaggcaaacttgaggcgccacacccgaacacatacaggagaaaagccattccaatgtcgaatttgtatgcgcaatttttcagatcactcaagcctcaagcgacacctccgcacacatactggttcacagaagccctttcagtgcaggatttgtatgcgcaactttagccaatcagcgaaccttttgcggcacactagaacgcatacaggtgagaagcctttccagtgtcgcatctgtatgcggaacttcagcgaccccagttcattgaagaggcatttgcgaactcacaccggttctcaaaaaccttttcagtgccgaatttgtatgcgcaacttcagccaacaaacaaatttgacgagacacacgcgcacgcacaccggggaaaaaccgtttcaatgtcgaatctgtatgcgcaattttagcgatgcgacacaacttgttaggcatctgcgcacacacttgcgg | 528 | LSQLPGERPFQCRICMRNFSEEANLRRHTRTHTGEKPFQCRICMRNFSDHSSLKRHLRTHTGSQKPFQCRICMRNFSQSANLLRHTRTHTGEKPFQCRICMRNFSDPSSLKRHLRTHTGSQKPFQCRICMRNFSQQTNLTRHTRTHTGEKPFQCRICMRNFSDATQLVRHLRTHLR | 176 | cGTCGAAGTCGAAGTCGACc |

**Supplemental Table 1 |** DNA and protein sequence of all Zinc Fingers used in this study and their target sequences.

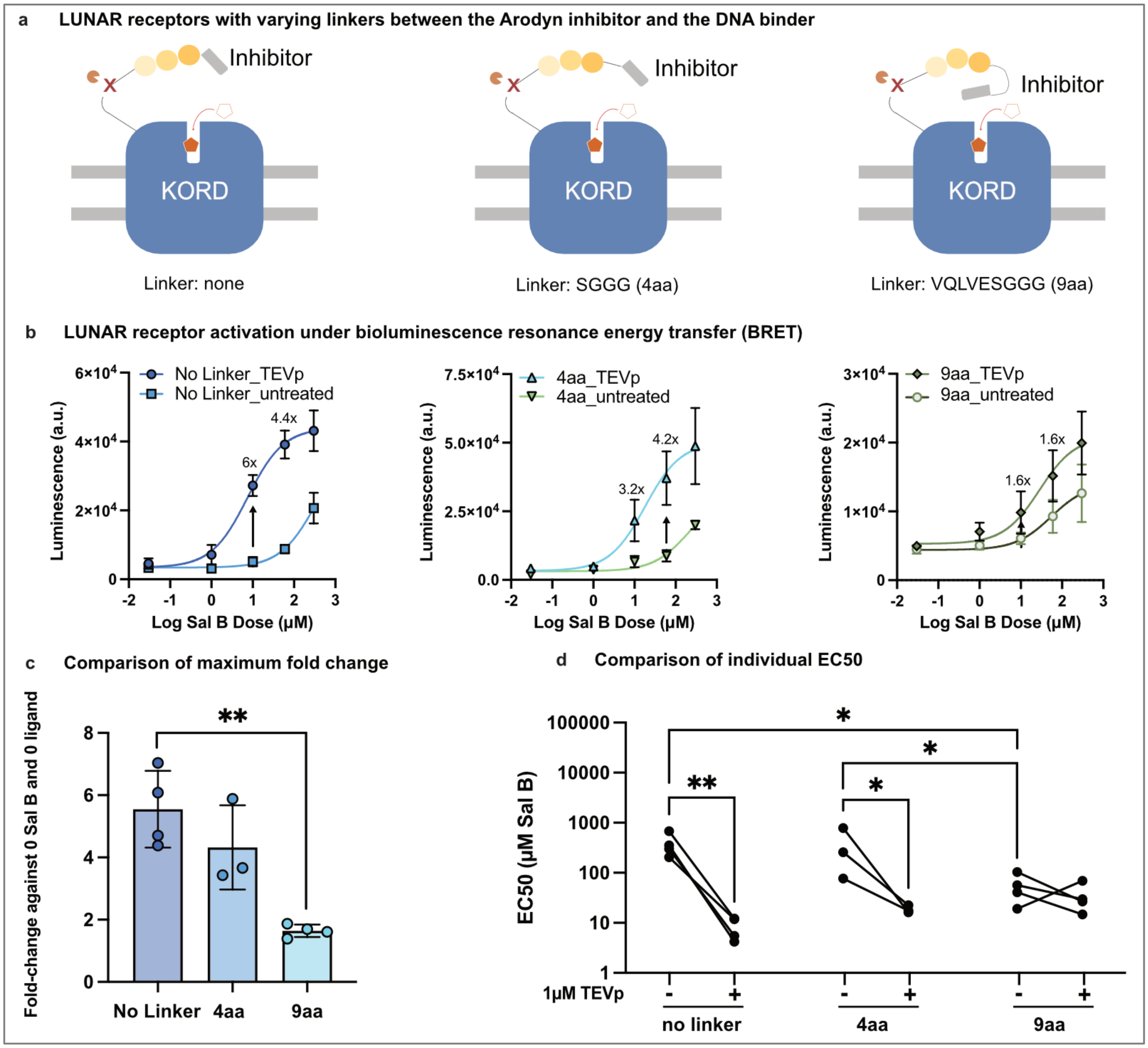


**Supplemental Fig. 1 | Linker scan and TEVp activation of Zinc Finger-based LUNAR. a.** Diagrams of a synthetic 3-finger LUNAR with no linker connecting the peptide inhibitor and the Zinc Finger DNA binding domain and its 4-amino acid (SGGG) and 9-amino acid (VQLVESGGG) linker variants. All variants were treated media without (-) and with (+) 1µM TEV Protease (TEVp), then activated by a gradient of Sal B and NanoLuc substrate. **b.** The Sal B dose response curves of three variants with different range of activation. **c.** Comparison of maximum fold change across 3 linker variant: for the no linker version, the maximum fold change from individual biological replicates is 5.5 ± 1.2 fold at 10 µM Sal B, for the 4aa variant, the maximum fold change is 4.3 ± 1.4 fold at 60 µM Sal B, and for the 9aa variant, the maximum fold change is 1.6 ± 0.2 fold at 60 µM Sal B. **d.** Comparison of EC50s without and with TEVp treatment for the linker variants showing the EC50 shift is the most significant for the no linker group and least significant for the longest linker group (n = 3-4 independent biological replicates, each with 2 technical replicates, *p<0.05, **p<0.01).

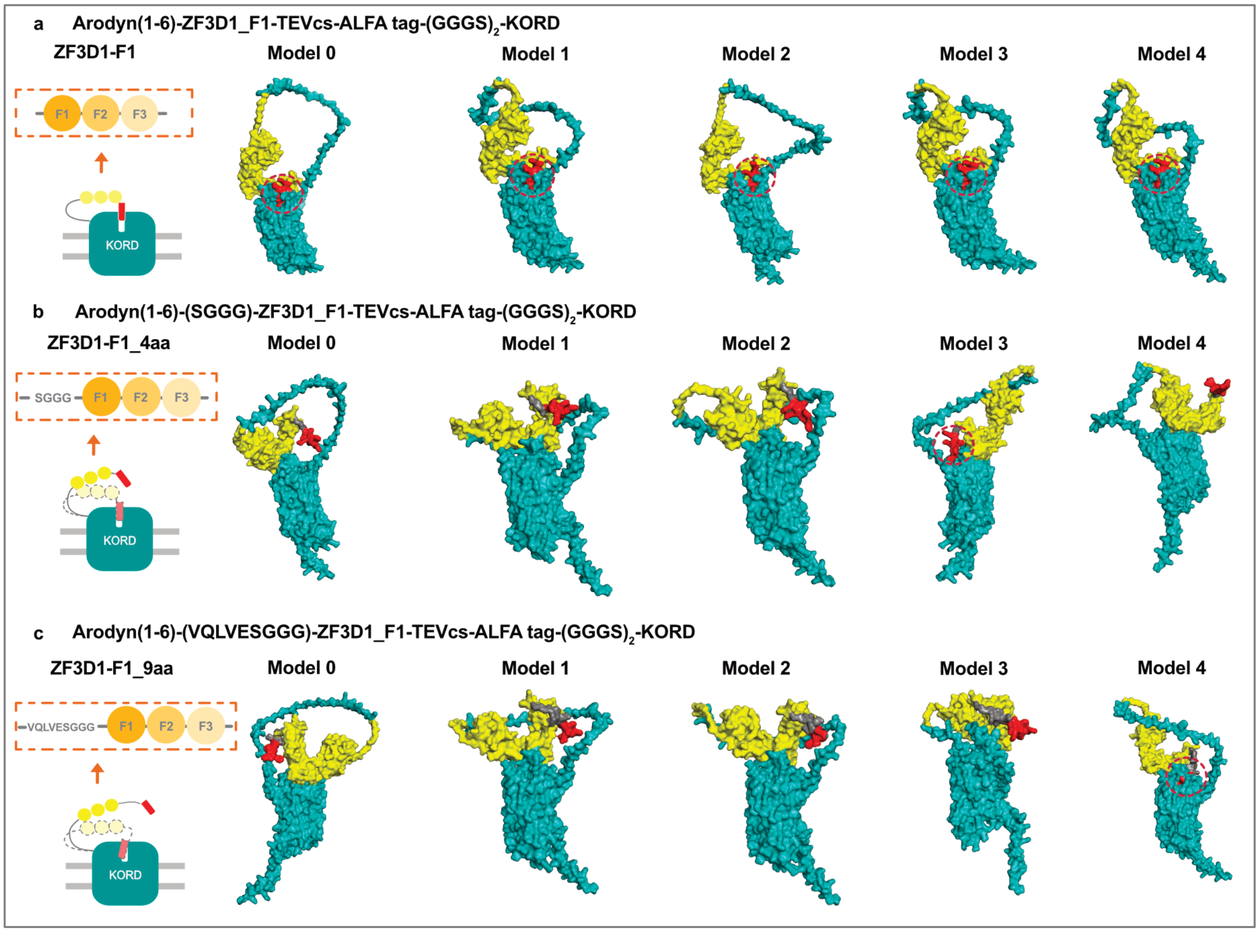


**Supplemental Fig. 2 | Structure Prediction of ZF3D1 receptors. a.** ZF3D1_F1 without any linkers, modeled in AF3 from the peptide inhibitor Arodyn (red), ZFP binder domain (yellow) to the transmembrane KORD (teal). **b.** ZF3D1 with a 4aa linker (gray) modeled in AF3, in same colors, and **c.** ZF3D1 with a 9aa linker (in gray) modeled in AF3. Dashed red circle show arodyn inhibitor sitting inside the orthosteric pocket of KORD, as shown in **a**-model 0, 1, 2, 3, 4; **b**-model 3; and **c**-model 4.

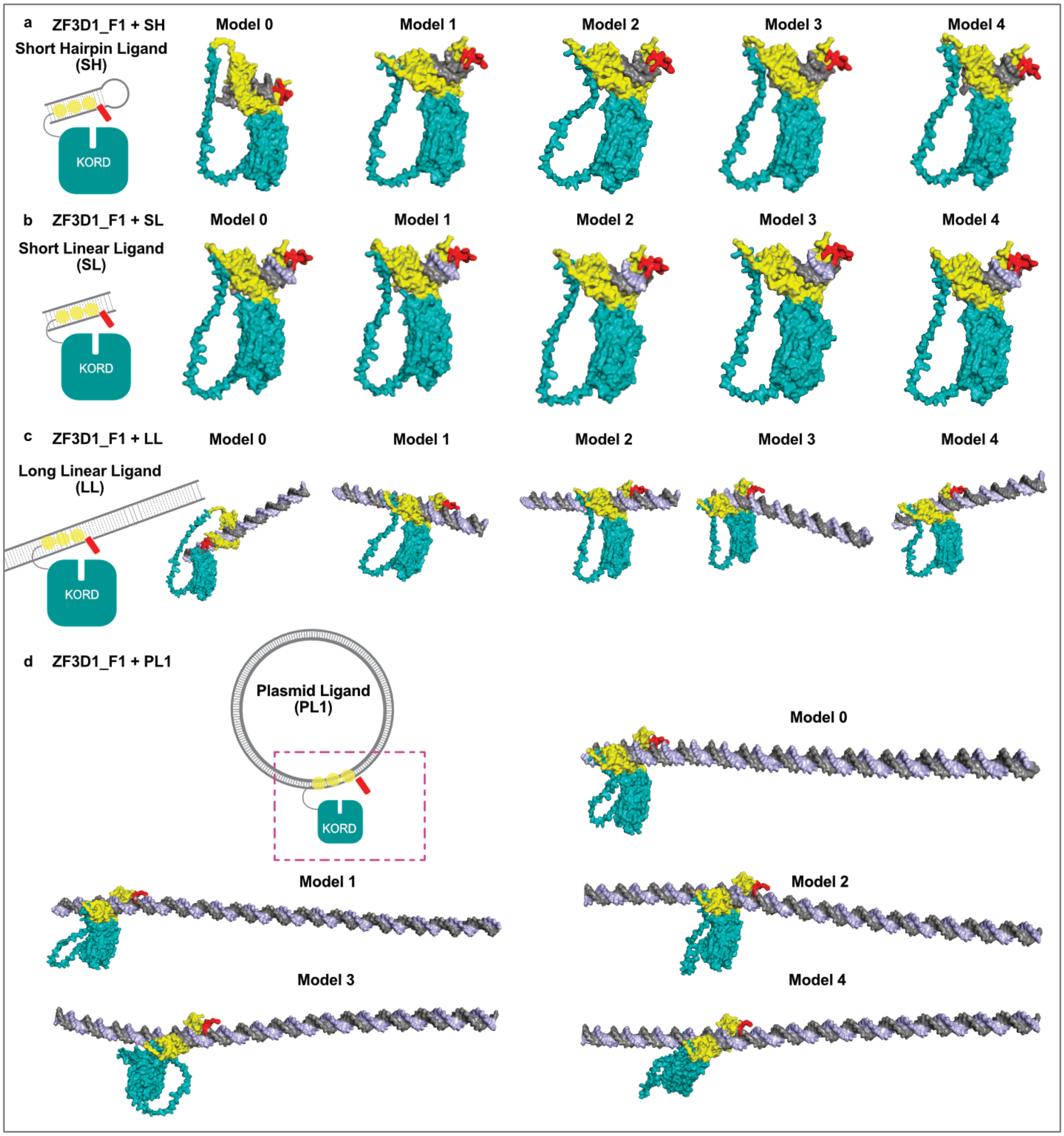


**Supplemental Fig. 3 | Structure Prediction of ZF3D1_F1 and various DNA ligands. a.** ZF3D1_F1 with short hairpin DNA ligand SH. **b.** ZF3D1 with short linear ligand SL, **c.** ZF3D1 with long linear ligand LL (contains 4 binding targets), and **d.** ZF3D1 with a 135bp section of the Plasmid ligand PL1 (in magenta rectangle with dashed lines, contains 4 binding targets).

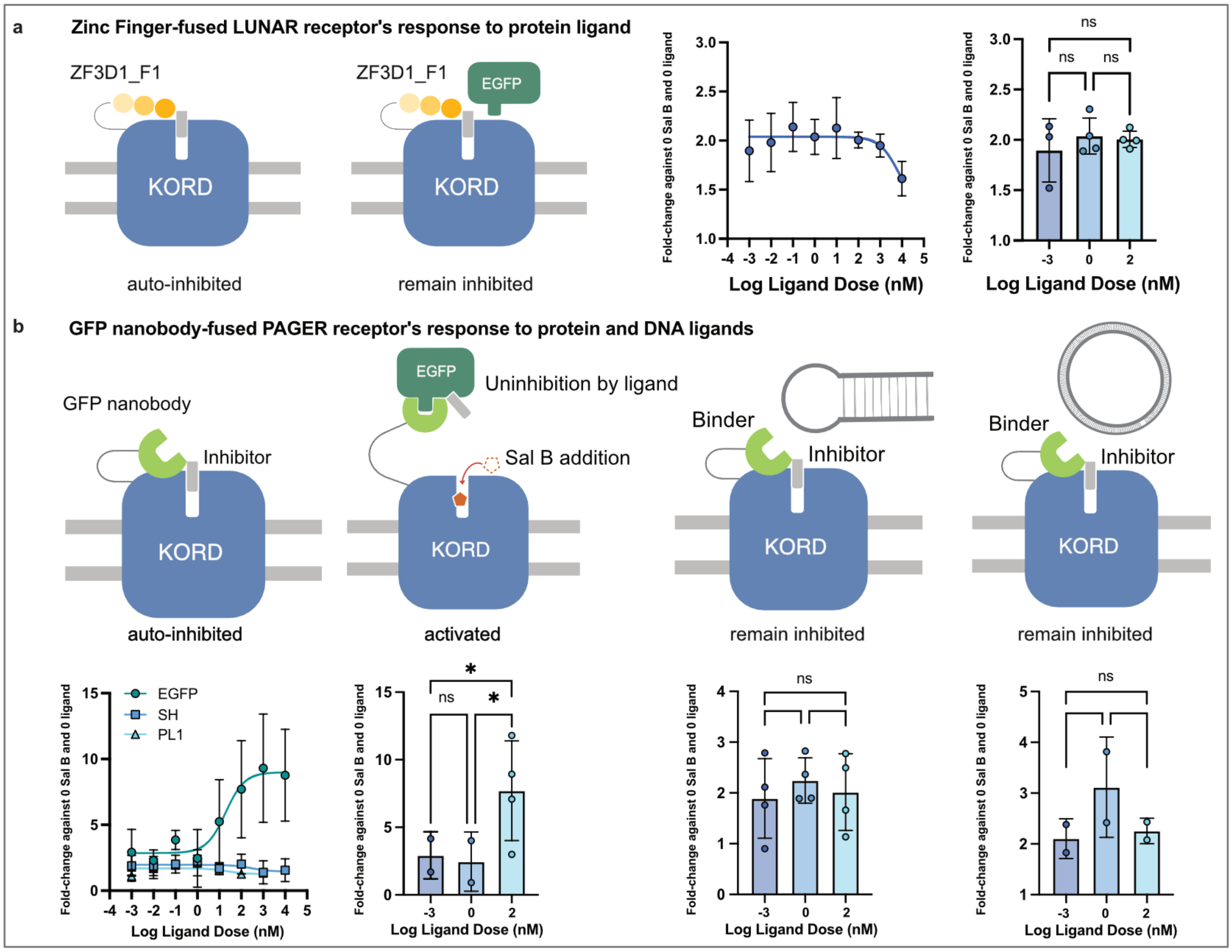


**Supplemental Fig. 4 | LUNAR activation is molecular specific. a.** 3-finger ZF LUNAR receptor ZF3D1-F1 remains inhibited upon addition of gradient EGFP as no significance of fold activation was observed at selected ligand concentrations **b.** DNA Ligands cannot be detected by GFP nanobody PAGER, as shown in the gradient activation line graph. GFPnb PAGER can be activated upon addition of gradient EGFP while remaining inhibited upon addition of short hairpin ligand SH or plasmid ligand PL1. (n = 2-4 biological replicates, each with 2 technical replicates. * = $p<0.05$, ns = not significant)

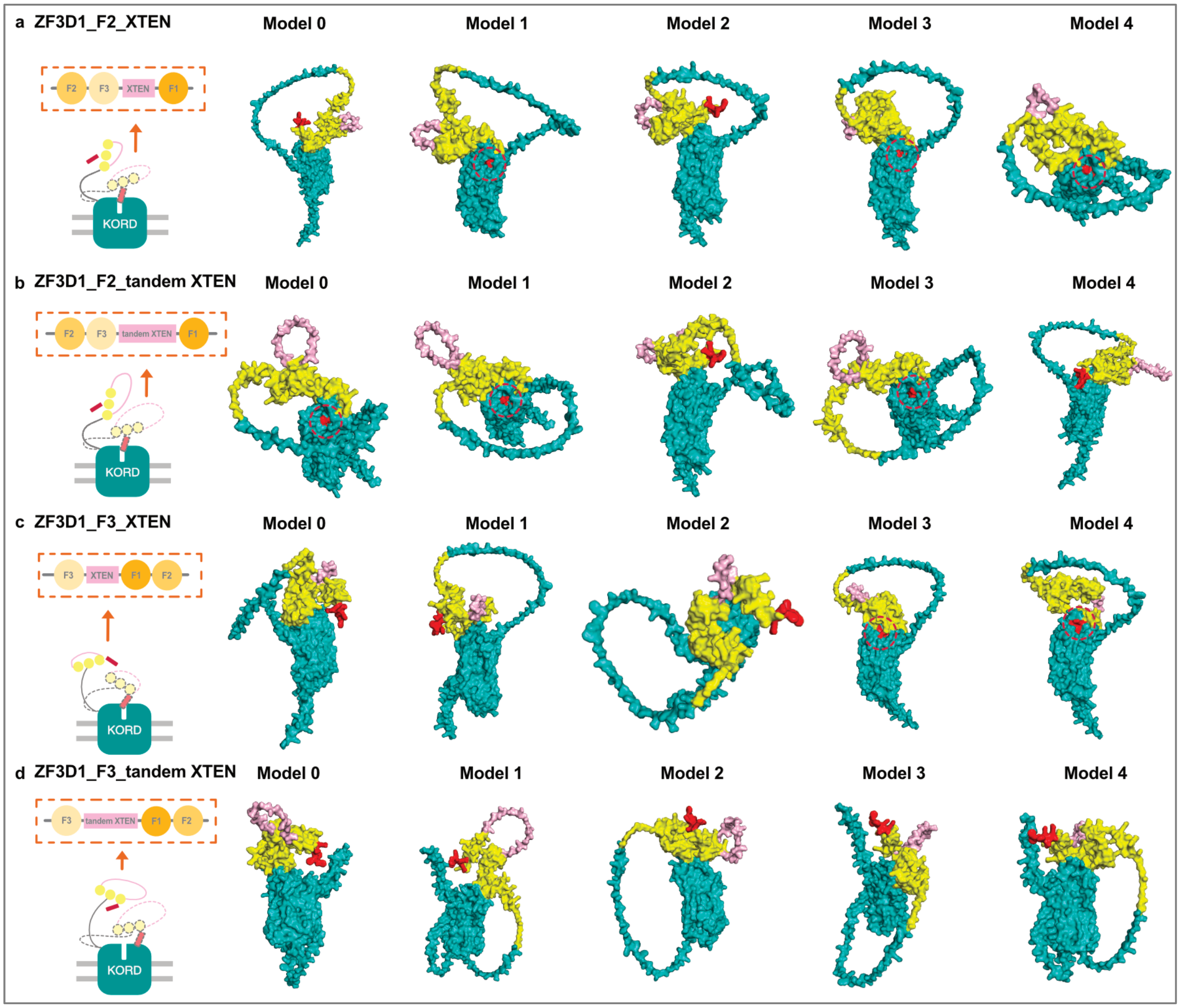


**Supplemental Fig. 5 | Structure Prediction of ZF3D1 circularly permutated receptors.** AlphaFold 3 modeling of permutants of ZF3D1, in the form of Arodyn (red), ZFP binder (yellow), and KORD (teal) containing **a.** ZF3D1_F2 binder (yellow) with one XTEN linker (pink), **b.** ZF3D1_F2 binder (yellow) with tandem XTEN linkers (pink), **c.** ZF3D1_F3 with one XTEN linker (pink), and **d.** ZF3D1_F3 with tandem XTEN linkers (pink). Dashed red circles show arodyn inhibitor sitting inside the orthosteric pocket of KORD, as shown in **a**-model 2, 3, 4; **b**-model 0, 1, 3; and **c**-model 3, 4.

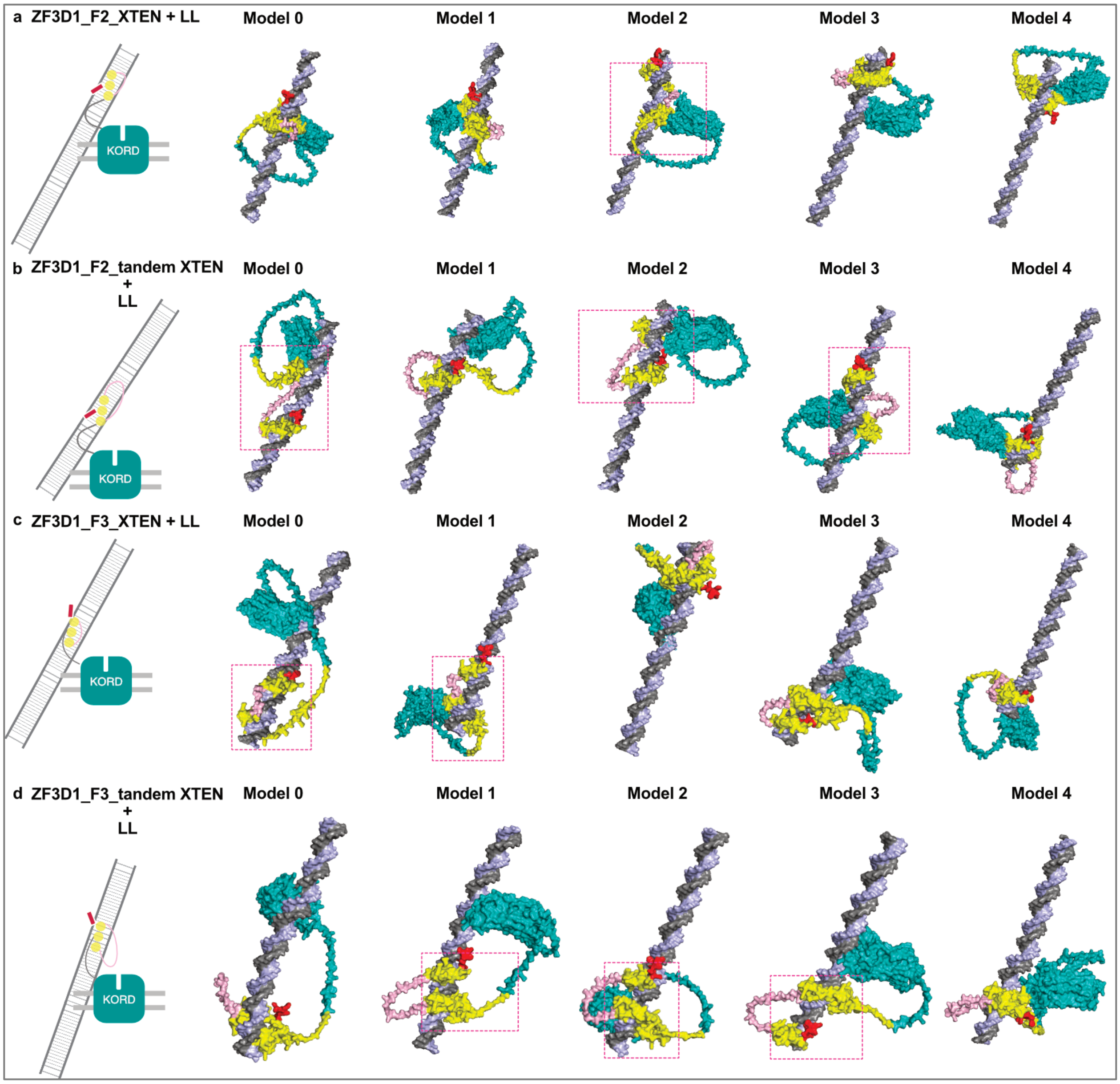


**Supplemental Fig. 6 | Structure Prediction of ZF3D1 permutants with long linear DNA ligand (LL).** AlphaFold 3 modeling of permutants of ZF3D1, in the form of Arodyn (red), ZFP binder (yellow), and KORD (teal) containing **a.** ZF3D1_F2 binder (yellow) with one XTEN linker (pink), **b.** ZF3D1_F2 binder (yellow) with tandem XTEN linkers (pink), **c.** ZF3D1_F3 with one XTEN linker (pink), and **d)** ZF3D1_F3 with tandem XTEN linkers (pink). Magenta square with dashed lines indicate poorly folded Zinc Fingers, as shown in **a**-model 2; **b**-model 0, 2, 3; **c**-model 0, 1; and **d**-model 1, 2, 3.

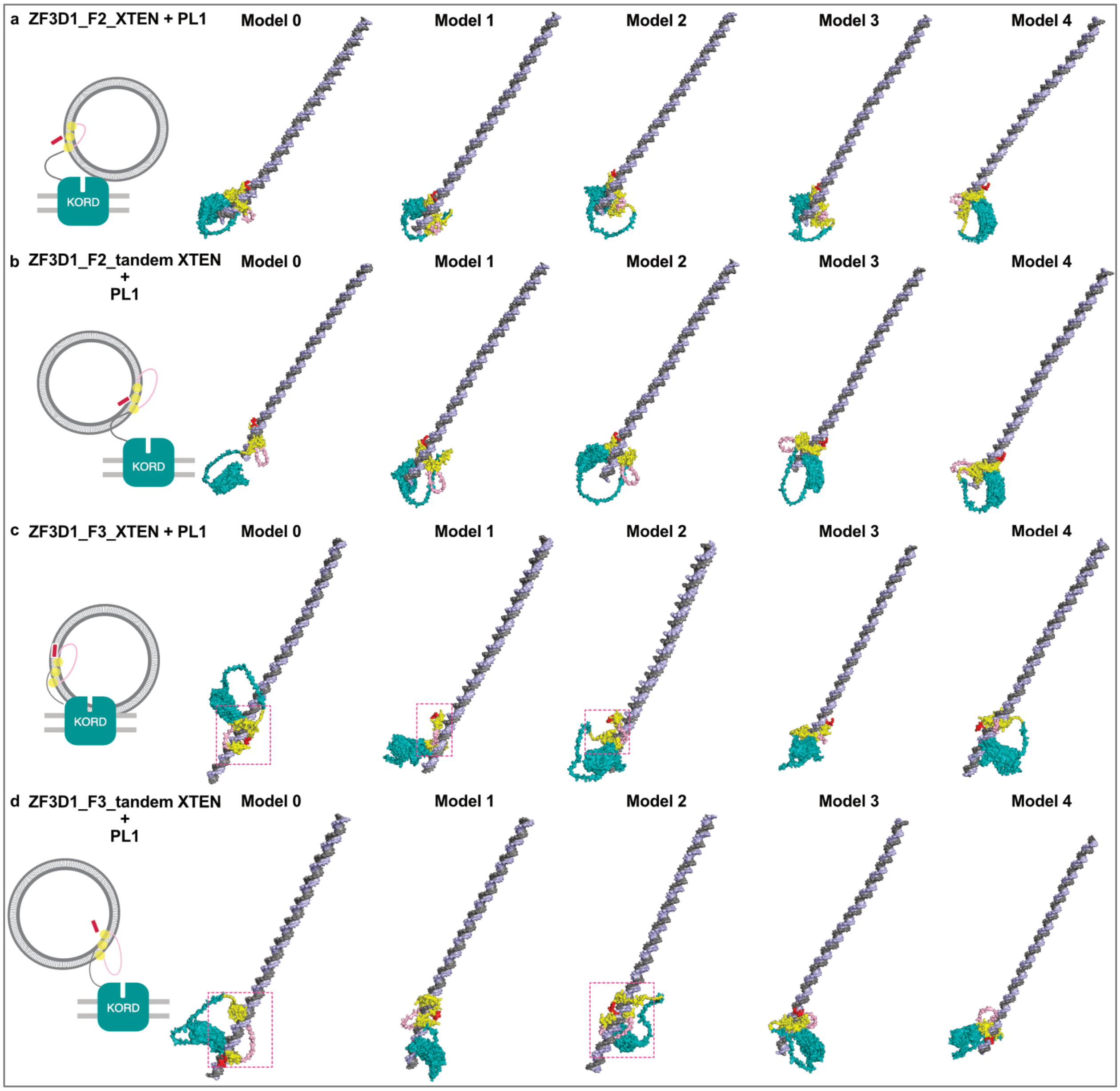


**Supplemental Fig. 7 | Structure Prediction of ZF3D1 permutants with an 135bp section of the plasmid DNA ligand (PL1).** AlphaFold 3 modeling of permutants of ZF3D1, in the form of Arodyn (red), ZFP binder (yellow), and KORD (teal) containing **a.** ZF3D1_F2 binder (yellow) with one XTEN linker (pink), **b**. ZF3D1_F2 binder (yellow) with tandem XTEN linkers (pink), **c.** ZF3D1_F3 with one XTEN linker (pink), and **d.** ZF3D1_F3 with tandem XTEN linkers (pink). Magenta rectangles with dashed lines indicate poorly folded Zinc Fingers, some of which resulted in individual fingers not latching onto the major groove of dsDNA, as shown in **c**-model 0, 1, 2; and **d**-model 0, 2.

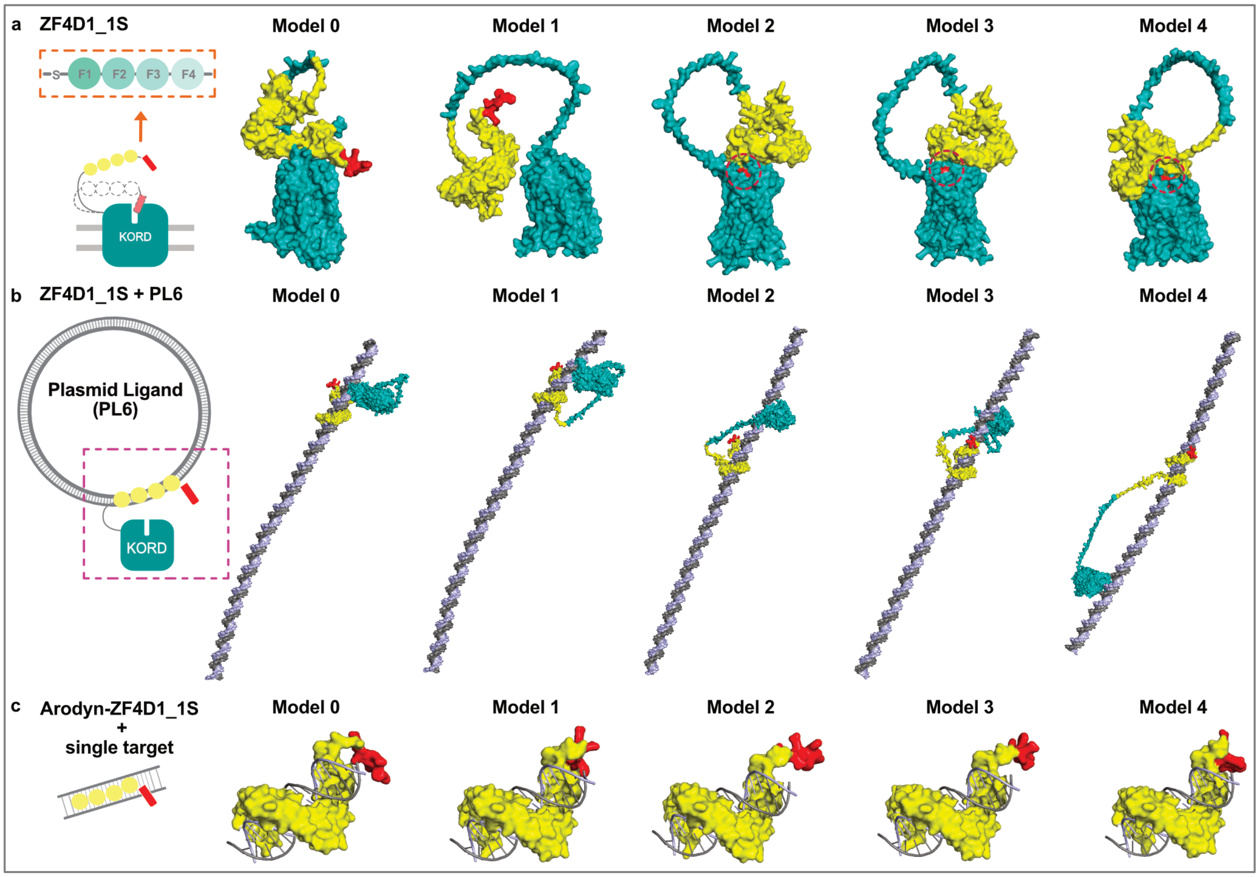


**Supplemental Fig. 8 | Structure Prediction of ZF4D1_1S receptor and matching ligand (PL6).** AlphaFold 3 modeling of **a.** the extracellular-transmembrane part of the ZF4D1_1S receptor in the form of Arodyn (red), ZFP binder (yellow), KORD (teal), **b.** ZF4D1_1S binding with an 165bp segment of the matching plasmid ligand PL6 (in magenta rectangle with dashed lines, contains 4 binding targets), **c.** ZF4D1_1S binding domain latching onto the major groove of a single dsDNA target. Dashed red circles show arodyn inhibitor sitting inside the orthosteric pocket of KORD, as shown in **a**-model 2, 3, 4.

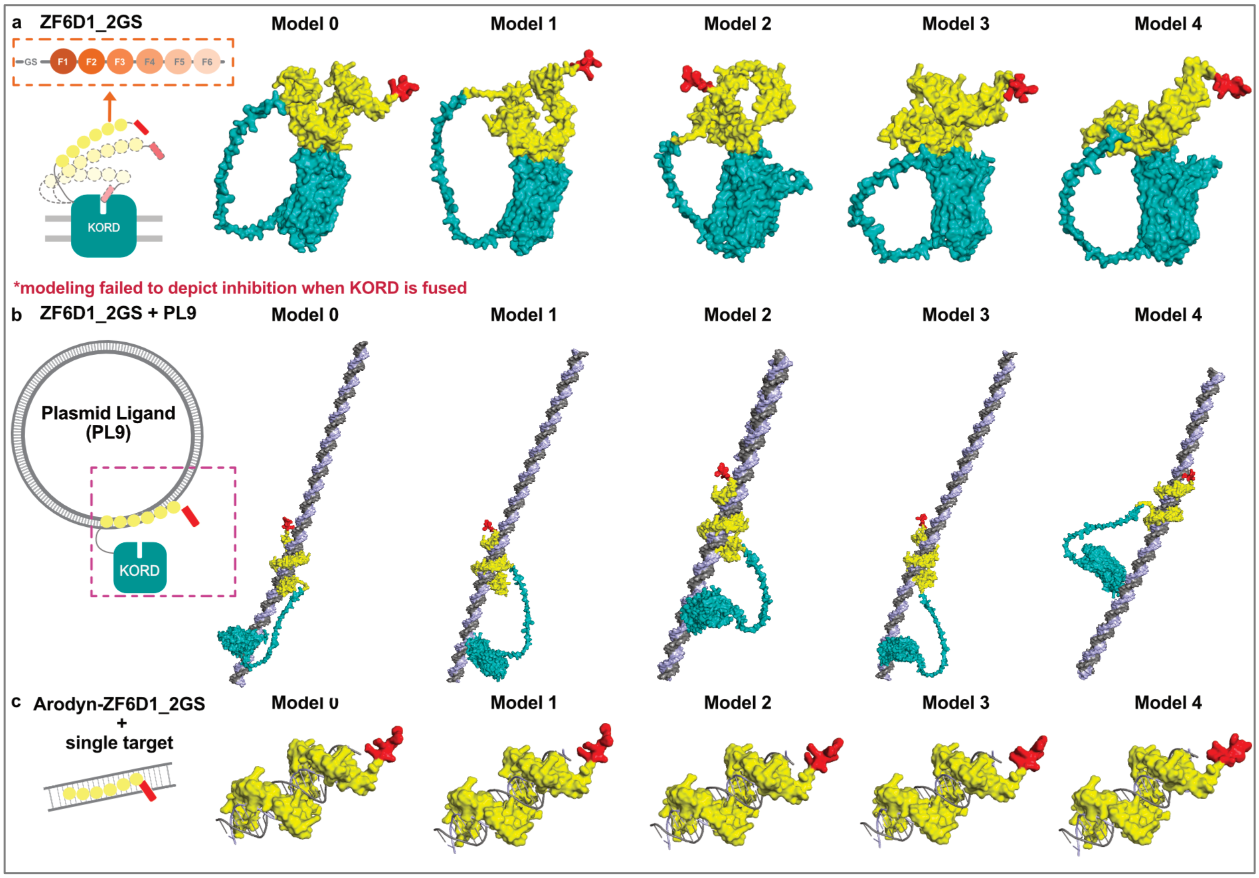


**Supplemental Fig. 9 | Structure Prediction of ZF6D1_2GS receptor and matching ligand (PL9).** AlphaFold 3 modeling of **a.** the extracellular-transmembrane part of the ZF6D1_2GS receptor in the form of Arodyn (red), ZFP binder (yellow), KORD (teal), **b.** ZF6D1_2GS binding with an 140bp segment of the matching plasmid ligand PL9 (in magenta rectangle with dashed lines, contains 4 binding targets), and **c.** ZF6D1_2GS binding domain placed within the major groove of a single dsDNA target.

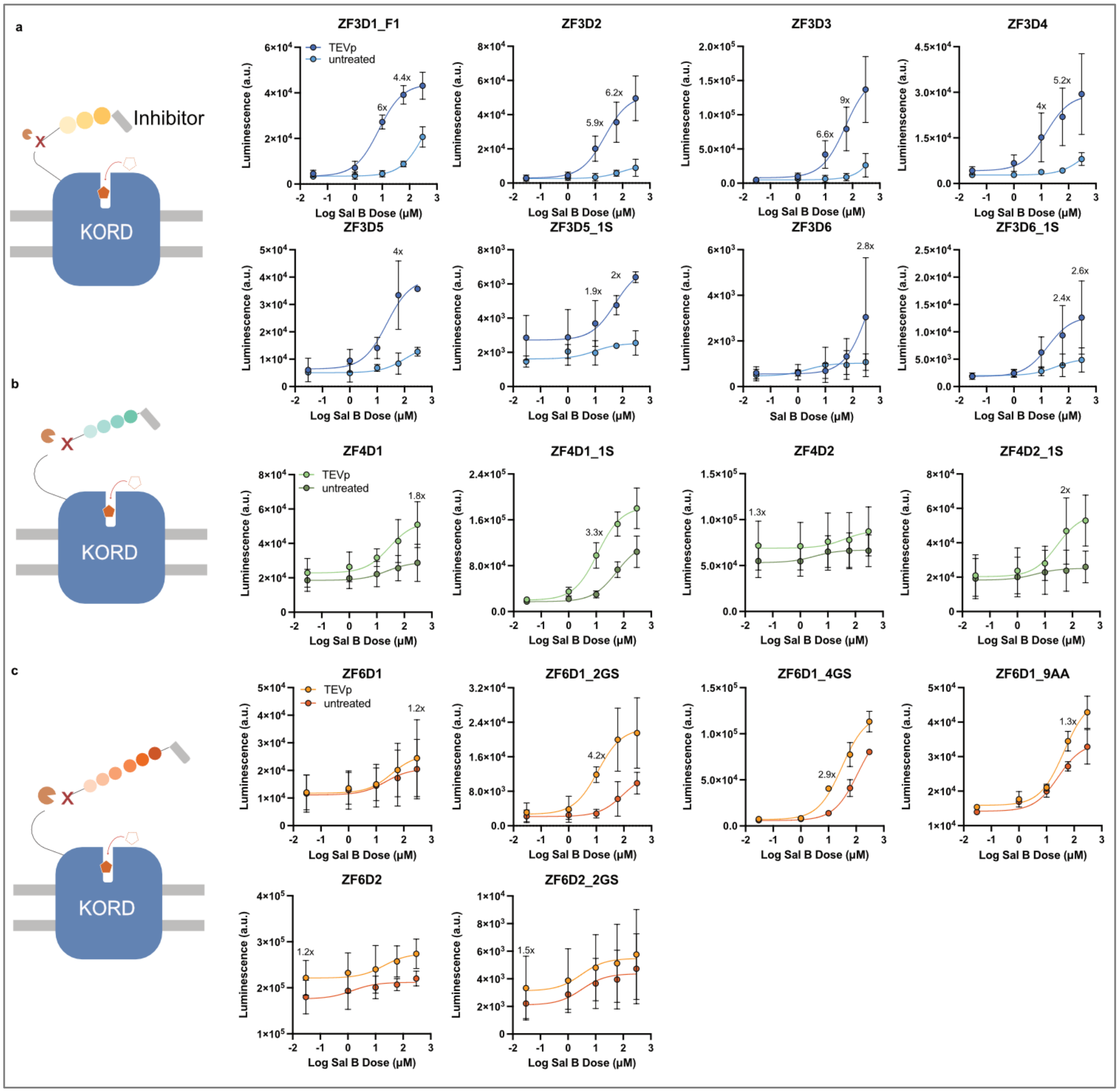


**Supplemental Fig. 10 | Sal B activation of all LUNAR constructs.** Diagram of inhibitor removal and LUNARs showing dose dependent activation by Sal B via inhibitor removal, including **a.** 3-finger (blue lines), **b.** 4-finger (green lines), **c.** 6-finger (orange lines) variants, although some binders required linker optimization (n = 2-6 biological replicates, each with 2 technical replicates).

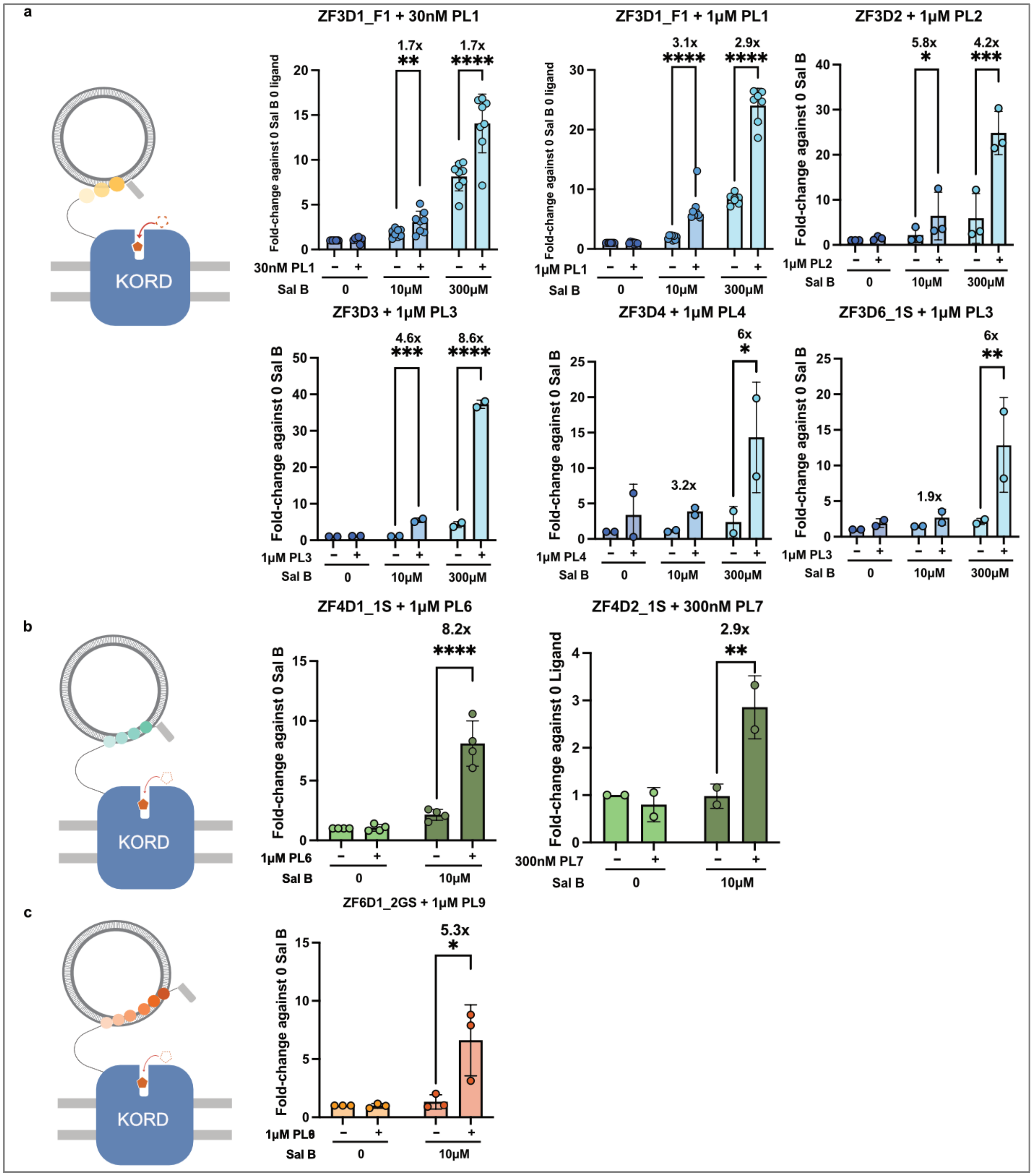


**Supplemental Fig. 11 | Plasmid DNA activation of LUNARs.** Zinc finger-based LUNAR receptors with **a.** 3-finger (blue bars), **b.** 4-finger (green bars), and **c.** 6-finger (orange bars) binders consistently detect plasmid DNA at various concentrations with a significant fold-change from the untreated group at the same Sal B concentration (n = 2-8 biological replicates, each with 2 technical replicates * = p<0.05, **p<0.01, ***p<0.001, ****p<0.0001, 2-way ANOVA).

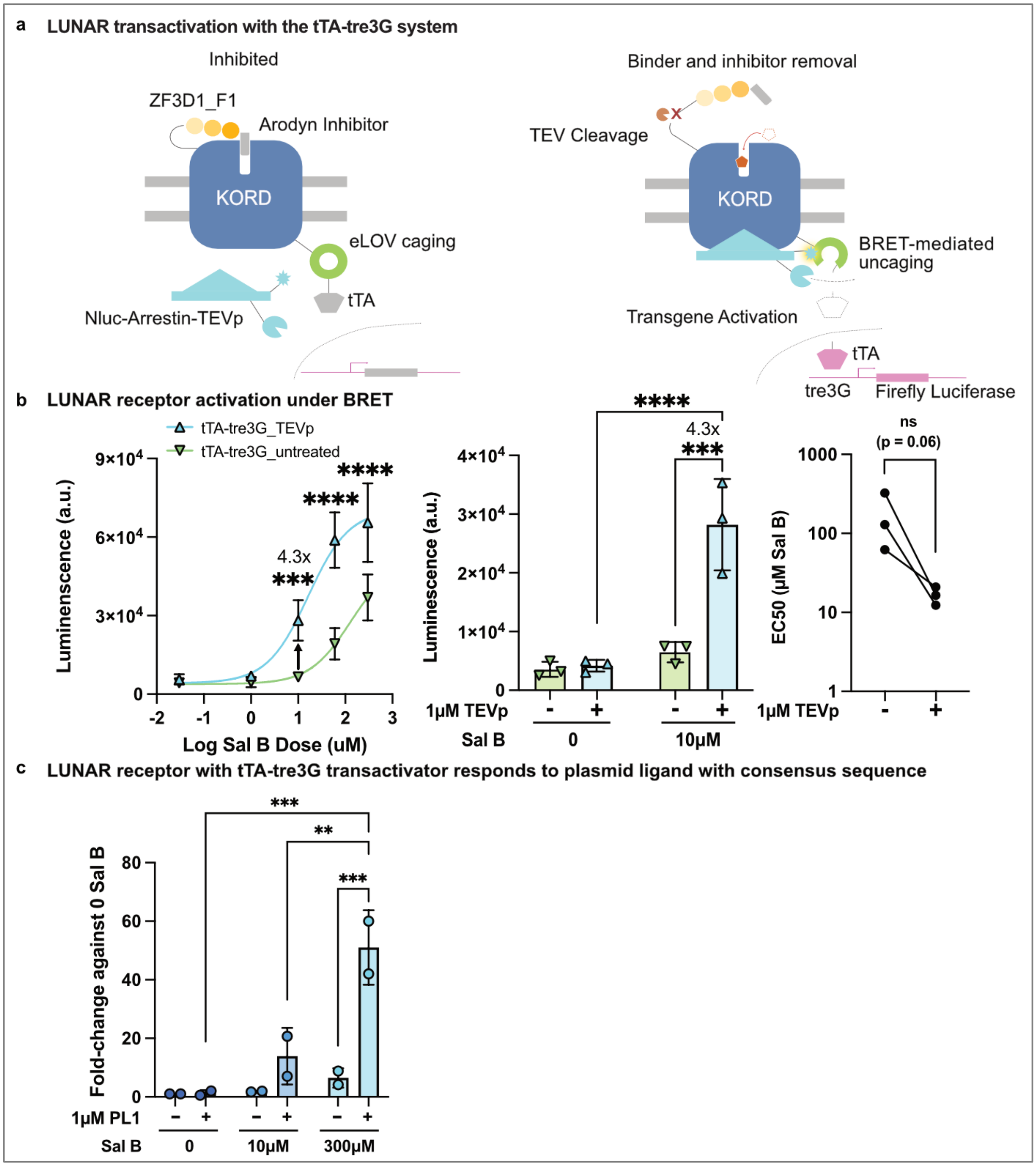


**Supplemental Fig. 12 | Characterization of a LUNAR receptor with a different transactivator.** **a.** Diagram of ZF3D1_F1 synthetic LUNAR receptors with the tTA2-tre3G transactivation circuit. **b.** Characterization of receptor responses to Sal B without (-) and with (+) TEV protease (TEVp) treatment under the tTA2-tre3G transactivator system, the bar charts featuring responses at 0 µM and 10 µM Sal B, and EC50 changes. **c.** Activation of ZF3D1_F1 with circular plasmid ligand PL1 and at 30 nM, 1 µM, or 10 µM ligand concentrations. (n = 2-3 biological replicates, each with 2 technical replicates **p<0.01, ***p<0.001, ****p<0.0001, ns = not significant, 2-way ANOVA).
